# Ribophorin-1 Governs Spike Abundance of Highly Pathogenic Coronaviruses by ER-associated degradation

**DOI:** 10.64898/2026.05.20.726704

**Authors:** Yi-Jiao Huang, Lu Lv, Wei Ran, Hui Zhao, Yong-Qiang Deng, Qing Ye, Lin Li, Xiao-Yan Wu, Qi Chen, Yi Hu, Bei Sun, Jian-Jun Zhao, Jincun Zhao, Cheng-Feng Qin

## Abstract

The spike protein of highly pathogenic coronaviruses is indispensable for viral entry, pathogenesis, and immune evasion. However, specific host factors governing spike protein degradation and abundance on progeny virions remain largely uncharacterized. Here, we identify Ribophorin-1 (RPN1), a non-catalytic subunit of the oligosaccharyltransferase (OST) complex, as an ER-localized host restriction factor that selectively depletes spike to inhibit coronavirus infection. Conditional knockout of Rpn1 in mouse lung markedly exacerbates SARS-CoV-2 pathogenesis, increasing viral load, immune cell infiltration, and syncytium formation. In cells, RPN1 knockdown enhances susceptibility to diverse variants, whereas its overexpression attenuates infection. Nano-flow cytometry, cryo-EM and LSCM show that RPN1 reduces spike abundance on progeny virions, resulting in reduction in subsequent syncytium formation across SARS-CoV, SARS-CoV-2, and MERS-CoV. Mechanistically, RPN1 promotes VCP/p97-dependent retrotranslocation and AMFR-mediated ubiquitination of spike protein. Heterologous expression of RPN1’s functional domain (residues 180-307) confers significant protection against SARS-CoV-2 both *in vitro* and *in vivo*. Collectively, our findings define an RPN1-initiated ERAD pathway that selectively targets coronavirus spike for proteasomal destruction, and provide proof-of-concept evidence that targeting this RPN1-dependent pathway represents a promising broad-spectrum antiviral strategy.

## Introduction

Three highly pathogenic coronaviruses—SARS-CoV-2, SARS-CoV and MERS-CoV—pose a persistent threat to global public health^1^. Although current clinically advanced antiviral strategies primarily target viral proteins — specifically, viral proteases essential for replication or the spike (S) protein responsible for host receptor binding—specific host-directed strategies targeting coronavirus structural protein quality control remain relatively underexplored^2^. Complementarily, host-directed therapies have emerged as a promising approach to evade viral resistance, as proteomics- or CRISPR-based screening has identified numerous host factors that regulate the viral life cycle^3–5^. Nevertheless, ER-localized host restriction factors—particularly those acting through ER-associated degradation (ERAD) — remain poorly characterized, and their therapeutic potential as druggable targets has not been experimentally evaluated.

SARS-CoV-2, SARS-CoV, and MERS-CoV each encode three structural glycoproteins—spike, membrane (M), and envelope (E)—embedded in the viral envelope^6,7^. During coronavirus replication, these structural proteins are synthesized, folded, and trafficked primarily through the ER^8,9^. Among them, the spike protein is a coronavirus-specific hallmark protein that forms the characteristic crown-like morphology surrounding the virion and functions as the “molecular key” for host cell entry, mediating both receptor binding and membrane fusion^10–12^. It is not only the principal target for vaccine development and neutralizing antibody therapeutics but also a driver of pathological syncytium formation and long COVID^13–15^. Structural biology study have revealed that individual SARS-CoV-2 virions display a random distribution of 24 ± 9 spike trimers, with this number varying across different viral variants^16^. The abundance of spike protein on the virion surface is a critical determinant of viral infectivity; naturally occurring mutations (e.g., D614G and Omicron sublineages of SARS-CoV-2) that alter spike protein expression or incorporation into virions directly influence viral pathogenicity and transmission^6,17–20^. However, the intrinsic mechanism by which host factor govern the intracellular biogenesis and degradation dynamics of the coronavirus spike protein, as well as the spike abundance on progeny virions remains largely unknown.

Ribophorin-1 (RPN1) is a core structural subunit of oligosaccharyltransferase (OST) complex, anchoring them and facilitating co-translational N-glycosylation at the ribosome–SEC61 translocon interface^21,22^. RPN1 functions as a substrate-specific adaptor that selectively engages nascent polypeptides bearing glycosylation-competent sequons and delivers them to the catalytic STT3 subunits of the OST complex—without possessing intrinsic glycosyltransferase activity^23^. Structurally, RPN1’s cytosolic domain supports assembly of the ribosome-translocon-OST-TRAP supercomplex by mediating TRAP recruitment to the ribosome, whereas its luminal domain serves as a central scaffold for OST subunits and ensures precise positioning of nascent polypeptides relative to the catalytic center for efficient co-translational N-glycosylation^22,24^. Beyond its canonical role in glycosylation, RPN1 has been implicated in ER-phagy, chaperone-mediated recognition of misfolded proteins, and cancer progression^25–27^. To date, no direct role for RPN1 in the viral life cycle—beyond its essential function in viral glycoprotein N-glycosylation — has been experimentally established.

In this study, we uncover an unanticipated role for RPN1 as an ERAD factor that selectively targets the spike protein of highly pathogenic coronaviruses, and heterologous expression of RPN1’s functional domain prevents viral infection *in vitro* and *in vivo*, establishing RPN1 as a promising broad-spectrum antiviral target.

## Results

### RPN1 deficiency enhances SARS-CoV-2 infection both *in vivo* and *in vitro*

Previously, we demonstrated that catalytic STT3 subunits of the OST complex are necessary for SARS-CoV-2 infectivity by mediating N-glycosylation of the spike protein^28^. Herein, to define the physiological role of RPN1 in coronavirus pathogenesis, we generated a lung epithelium–specific conditional knockout (cKO) mouse model (*Rpn1^fl/fl^, S*ftpc*^CreERT^*^2^) (Fig. 1a). Tamoxifen administration induced efficient deletion of Rpn1 in alveolar epithelial cells, as confirmed by immunoblotting and immunohistochemistry (Extended Data Fig. 1). Mice were then infected intranasally with the SARS-CoV-2 Beta variant, and analyzed at 3 days post-infection (dpi) (Fig. 1b). Given RPN1’s canonical role in OST-mediated N-glycosylation, which facilitates folding and trafficking of viral glycoproteins, Rpn1 deficiency was expected to attenuate infection. Surprisingly, *Rpn1*-cKO mice exhibited significantly elevated infectious viral titers in lung tissues (∼200-fold higher than wild-type controls), as quantified by focus-forming units (FFU), along with a ∼6-fold increase in viral RNA loads (Fig. 1c). Concurrently, *Rpn1*-cKO mice developed pronounced viremia, as evidenced by markedly elevated viral RNA levels in serum, whereas wild-type mice showed only minimal viral RNA (Fig. 1d). Western blot analysis confirmed both efficient depletion of endogenous Rpn1 in lung tissues and robust accumulation of viral N protein (Fig. 1e). Immunofluorescence (IF) analysis revealed markedly widespread and intense SARS-CoV-2 viral protein staining in the lungs of *Rpn1*-cKO mice, in stark contrast to the sparse, focal signal observed in wild-type controls (Fig. 1f). Consistent with these findings, histopathological assessment showed significantly exacerbated lung injury in *Rpn1*-cKO mice, characterized by inducible bronchus-associated lymphoid tissue (iBALT)-like peribronchiolar and interstitial lymphoid aggregates and extensive syncytium formation within the alveolar parenchyma—pathological features that were minimal or absent in wild-type mice (Fig. 1g,h, and Extended Data Fig. 2). These findings demonstrate that RPN1 exerts a non-redundant, host-protective function *in vivo* that counterbalances its proviral glycosylation activity.

**Fig. 1.**
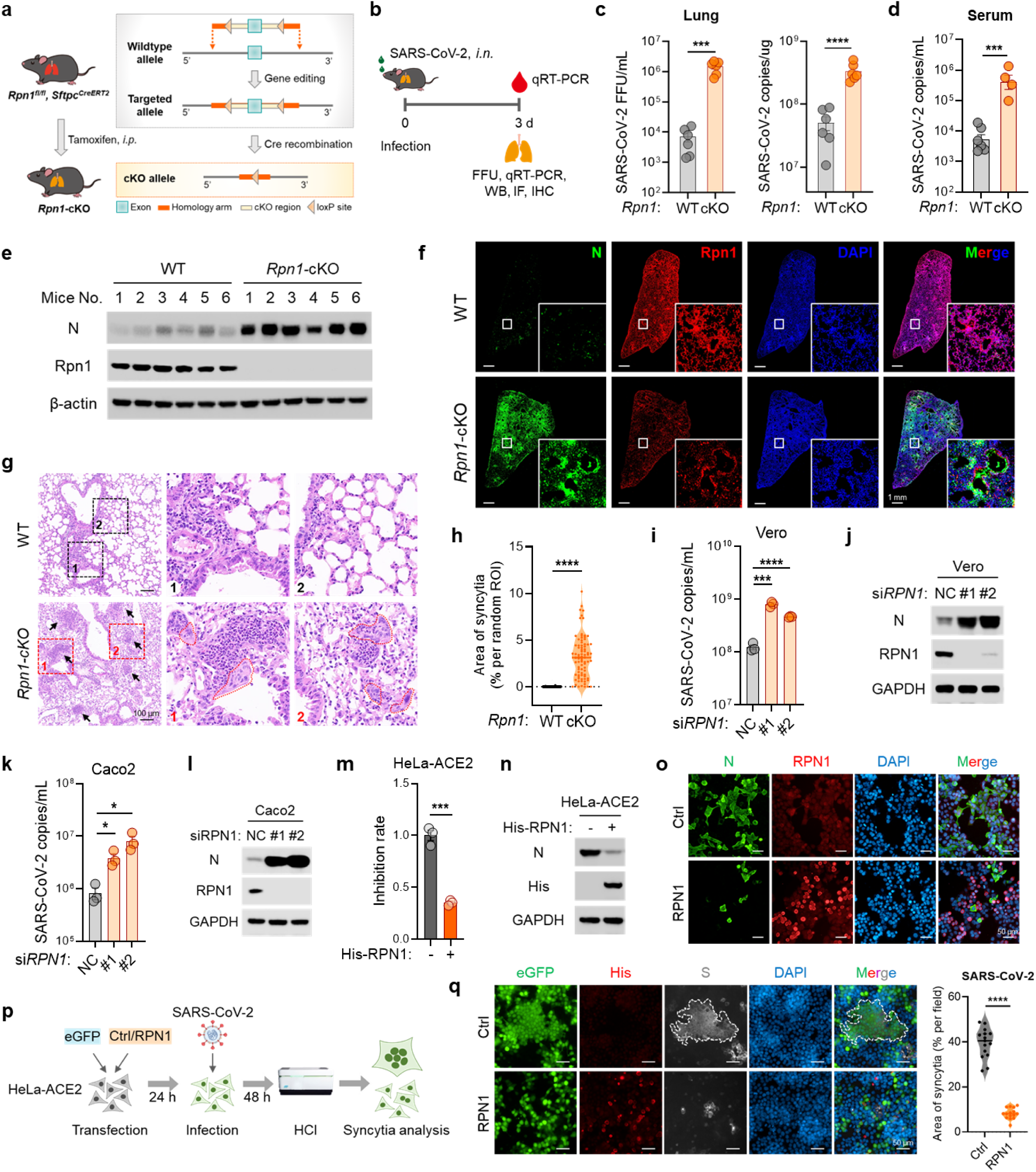
RPN1 inhibited SARS-CoV-2 infection both *in vivo* and *in vitro*. **a**, Strategy for generating conditional *Rpn1* knockout (*Rpn1*-cKO) mice. *Rpn1^fl/fl^, S*ftpc*^CreERT2^* mice were treated with tamoxifen intraperitoneally (*i.p.*) to induce deletion of Rpn1 specifically in alveolar epithelial cells. **b**, Schematic diagram of SARS-CoV-2 infection in mice. **c**-**h**, 7-month-old wild-type (WT) or *Rpn1*-cKO mice (n=6) were infected intranasally with SARS-CoV-2 Beta strain (1×10^4^ FFU). Lungs and serum were harvested at 3 days post-infection (dpi) for further analysis. (**c**) Viral loads in lungs: infectious virus titers quantified by focus-forming assay (left); viral RNA quantified by qRT-PCR (right). (**d**) Circulating viral RNA levels in serum, measured by qRT-PCR. (**e**) Immunoblot analysis of viral N protein and Rpn1 expression in lung lysates. (**f**) Immunofluorescence staining of lung sections; viral N protein (green), Rpn1 (red), and nuclei (DAPI, blue) are shown; boxed regions are enlarged in the bottom-right insets. (**g**) Representative H&E staining of lungs; arrows indicate focal lymphocytic aggregation; boxed regions are enlarged in the right; dashed lines in the enlarged boxes outline multinucleated syncytial structures. (**h**) Quantification of syncytial area in selected random region of interest (ROI) in lung sections. **i**,**j**, Vero cells were transfected with scrambled siRNA (negative-control, NC) or si*RPN1* for 48 h, then infected with SARS-CoV-2 (MOI = 0.01) for 48 h. Viral RNA in supernatants was quantified by qRT-PCR (n = 3) (**i**). Intracellular viral N protein and RPN1 levels were analyzed by immunoblotting (**j**). **k**,**l**, Caco2 cells were transfected with scrambled siRNA or si*RPN1* for 48 h, then infected with SARS-CoV-2 (MOI = 0.01) for 72 h. Viral RNA in supernatants was quantified by qRT-PCR (n = 3) (K). Intracellular viral N protein and RPN1 levels were analyzed by immunoblotting (L). **m**-**o**, HeLa-ACE2 cells were transfected with *His-RPN1* for 24 h prior to infection with SARS-CoV-2 (MOI = 0.01). (**m**) Viral RNA levels in supernatants, quantified by qRT-PCR, are presented relative to the control group (set to 1) to calculate the percent inhibition conferred by RPN1 group at 48 h post-infection (n = 3). (**n**) Intracellular viral N protein and His-RPN1 expression were analyzed by immunoblotting at 48 h. (**o**) Immunofluorescence staining showing viral N protein (green), RPN1 (red) and nuclei (DAPI, blue) at 48 h. **p**, Schematic diagram of the experimental workflow for SARS-CoV-2-induced syncytia analysis. **q**, eGFP was co-transfected with control or *His-RPN1* for 24 h prior to infection with SARS-CoV-2 delta strain (MOI = 0.01) in HeLa-ACE2 cells. Immunofluorescence staining was performed at 48 h post-infection to visualize syncytium formation. eGFP (green) was used to visualize SARS-CoV-2-induced membrane fusion. RPN1 (red), S protein (gray), and nuclei (DAPI, blue) are shown. Representative syncytia are marked with dashed borders. Quantitative analysis of syncytial area per microscopic field is presented on the right. β-actin or GAPDH serves as a loading control (**e**, **j**, **l**, and **n**). Data are presented as mean ± SEM of biological replicates, two-tailed unpaired *t*-test (**c**, **d**, **h**, **i**, **k**, **m**, and **q**). (* *P* < 0.05; ** *P* < 0.01; *** *P* < 0.001; **** *P* < 0.0001)

To validate this restriction activity at the cellular level, we performed loss- and gain-of-function experiments in multiple SARS-CoV-2-susceptible human and non-human primate cell models. Consistent with the *in vivo* findings, siRNA-mediated knockdown of RPN1 significantly increased SARS-CoV-2 replication in both Vero (African green monkey kidney) and Caco2 (human colorectal adenocarcinoma) cells, as quantified by increased viral RNA and N protein expression (Fig. 1i–l). Conversely, overexpression of human RPN1 in HeLa-ACE2 suppressed SARS-CoV-2 infection (Fig. 1m-o) and its progeny variants—including Beta, Delta, and Omicron JN.1 (Extended Data Fig. 3). Moreover, RPN1 overexpression significantly attenuated SARS-CoV-2-induced syncytium formation in HeLa-ACE2 cells (Fig. 1p,q). Collectively, these *in vivo* and *in vitro* data establish RPN1 as a host restriction factor that actively constrains SARS-CoV-2 infection.

### RPN1 reduced spike protein levels and abundance on viral particles

Given RPN1’s ER localization and established role in the biogenesis and quality control of membrane proteins, we systematically assessed its impact on all four SARS-CoV-2 structural proteins—S, E, M and N—all of which localize to or transit through the ER/Golgi system. Immunoblot analysis revealed that RPN1 overexpression significantly reduced spike protein levels, while exerting negligible effects on E, M, and N expression (Fig. 2a). This selectivity—combined with our prior observation that *Rpn1*-cKO mice exhibit a substantially greater increase in infectious viral titers (FFU) than in viral RNA loads (Fig. 1c), the established correlation between virion-associated spike protein abundance and infectivity^17,18^, and the exacerbated syncytium formation observed in lung tissues (Fig. 1g,h), a process strictly dependent on functional spike-mediated membrane fusion^14,29^—strongly implicates the spike protein as the primary molecular target through which RPN1 constrains coronavirus infection.

**Fig. 2.**
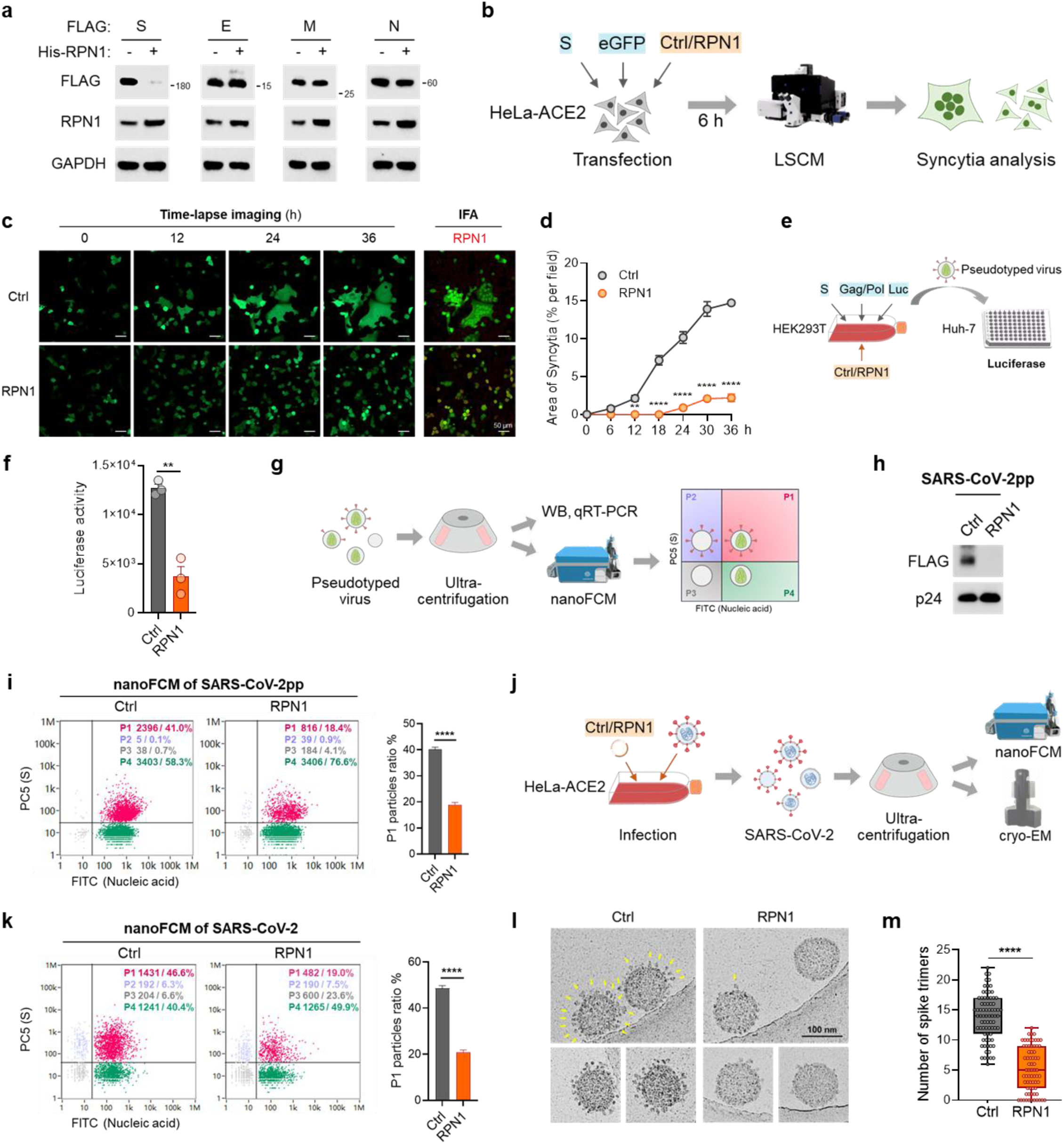
RPN1 reduced spike levels and abundance on the virus particles. **a**, Immunoblot analysis of FLAG-tagged SARS-CoV-2 spike (S), envelope (E), membrane (M), and nucleocapsid (N) protein in the presence or absence of RPN1 overexpression. GAPDH serves as a loading control. **b**, Schematic diagram of the experimental workflow for S-mediated syncytia analysis. **c**,**d**, HeLa-ACE2 cells were co-transfected with plasmids encoding S, eGFP, and control or RPN1. Live-cell time-lapse imaging was initiated 6 h post-transfection and continued for 36 h. See Supplementary Videos 1-2 for time-lapse movie. (**c**) Representative frames at 0, 12, 24, and 36 h into imaging show eGFP-labeled cell-cell fusion; after fixation at 37 h, immunofluorescence staining for RPN1 (red) is shown on the right. (**d**) Syncytial area per field was quantified every 6 h from time-lapse recordings. **e**, Schematic of SARS-CoV-2 pseudotyped virus production and infection assay. **f**, Pseudotyped virus were generated in HEK293T cells by co-transfection of plasmids encoding SARS-CoV-2 S, packaging protein (Gag/Pol), and a luciferase reporter (Luc) with control or RPN1. Huh7 cells were infected with pseudotyped virus preparations for 3 d, and luciferase activity was measured. (n = 3) **g**, Workflow for pseudotyped virus purification and analysis by immunoblotting, qRT-PCR and nanoscale flow cytometry (nanoFCM). **h**, Immunoblot analysis of FLAG-tagged S protein in purified SARS-CoV-2 pseudotyped particles (SARS-CoV-2pp) produced in control or RPN1-overexpressing cells. HIV Gag-p24 serves as a loading control. **i**, Single-particle analysis of purified SARS-CoV-2pp by nanoFCM. Representative density plots show SARS-CoV-2pp populations within the 80–150 nm size, stained for S protein (PC5-conjugated antibody) and nucleic acid (FITC). Four gates (P1–P4) were defined; P1 (red) corresponds to RNA⁺S⁺ intact particles. Percentages of events in each gate are indicated. Right: quantitative comparison of P1 particle frequency between control and RPN1-overexpressing conditions. **j**, Schematic of the authentic SARS-CoV-2 particle purification from infected HeLa-ACE2 cells followed by nanoFCM (**k**) and cryo-electron microscopy (cryo-EM) (**l** and **m**) analysis. **k**, NanoFCM analysis of purified SARS-CoV-2 virions. Representative density plots show SARS-CoV-2 virions populations within the 60–140 nm size. The frequency of P1 particles (RNA⁺S⁺) is quantified on the right. **l**,**m**, Cryo-EM analysis of purified SARS-CoV-2 virions. Yellow arrows indicate S protein on the viral surface (**l**). Quantification of S numbers per virion cross-section (**m**). Data are presented as mean ± SEM, two-tailed unpaired *t*-test (**d**, **f**, **i**, **k**, and **m**). (** *P* < 0.01; *** *P* < 0.001; **** *P* < 0.0001). All experiments (except for cryo-EM) were repeated at least three times independently.

To directly test whether RPN1 suppresses spike-dependent fusogenicity, we co-expressed spike protein, eGFP (indicating cell morphological) and RPN1 in HeLa-ACE2 cells—a model highly permissive for spike-mediated syncytium formation^30^. Live-cell laser scanning confocal microscopy (LSCM) demonstrated that expression of spike alone induced progressive, large-scale syncytia over 36 hours; in contrast, co-expression of RPN1 reduced the total syncytium area by >5-fold (Fig. 2b-d and Supplementary Videos 1-2). High-content image (HCI) analysis—using Na^+^/K^+^-ATPase immunostaining to delineate plasma membranes and DAPI staining to quantify nuclear content—confirmed these findings quantitatively (Extended Data Fig. 4). These data establish that RPN1 potently inhibits spike-driven cell–cell fusion.

Beyond syncytium formation, spike is indispensable for viral entry, because it must be properly incorporated into the envelope of progeny virions to mediate receptor binding and membrane fusion during the next round of infection^6,31^. To determine whether RPN1 compromises virion functionality via spike depletion, we employed two complementary systems, HIV-1–based spike-pseudotyped virus and authentic SARS-CoV-2. In the pseudotyped virus system, inclusion of an RPN1-expression plasmid in the standard three-plasmid packaging mix yielded pseudotyped particles (pp) with markedly reduced infectivity in Huh-7 cells (Fig. 2e,f), indicating impaired entry competence. Next, to systematically assess spike abundance on pseudovirions, we purified SARS-CoV-2 pseudotyped particles (SARS-CoV-2pp) by ultracentrifugation for subsequent biochemical and biophysical analysis (Fig. 2g). Western blot analysis revealed a significant and specific reduction in spike protein levels in SARS-CoV-2pp produced in RPN1-expressing cells, whereas viral genomic RNA and HIV Gag p24 (capsid protein) levels remained unchanged (Fig. 2h and Extended Data Fig. 5). Crucially, nanoscale flow cytometry (nanoFCM) of intact SARS-CoV-2pp confirmed that RPN1 overexpression reduced the proportion of spike- and RNA-positive particles (P1 gate: PC5^+^/FITC^+^) from ∼40% in control pseudovirions to ∼20%, thereby directly demonstrating diminished spike incorporation into virions (Fig. 2i). Furthermore, we purified authentic SARS-CoV-2 virions produced in RPN1-overexpressing HeLa-ACE2 cells to validate spike abundance on native viral particles (Fig. 2j). Consistent with pseudotyped virus results, nanoFCM analysis revealed that RPN1 overexpression significantly reduced the proportion of intact viral particles from ∼50% to ∼25% (Fig. 2k). Remarkably, Cryo-electron tomography (cryo-EM) of native SARS-CoV-2 virions confirmed this deficit at ultrastructural resolution: virions produced under RPN1 overexpression exhibited fewer spike trimers per cross section, averaging ∼15 trimers, compared with ∼5 in control virions (Fig. 2l,m). Collectively, these results demonstrate that RPN1 restricts SARS-CoV-2 propagation by reducing both the number of intact progeny virions and the spike abundance on individual viral particles.

### RPN1 induced the VCP/p97 complex-mediated ERAD of SARS-CoV-2 spike protein

To elucidate how RPN1 depletes spike protein, we first excluded transcriptional regulation. Quantitative RT–PCR confirmed that RPN1 overexpression did not alter *S* mRNA levels (Extended Data Fig. 6a). We therefore focused on post-translational mechanisms and assessed major proteolytic pathways. Treatment with the proteasome inhibitor MG-132—but not with lysosomal inhibitors, chloroquine (CQ), ammonium chloride (NH_4_Cl), bafilomycin A1 (baf-A1) or 3-methyladenine (3MA)—stabilized spike when inhibiting translation by cycloheximide (CHX) (Fig. 3a), indicating that spike is degraded primarily via the ubiquitin–proteasome system. Notably, MG-132 fully rescued the reduction of spike levels in RPN1-overexpressing cells, whereas chloroquine had no effect (Fig. 3b and Extended Data Fig. 6b), demonstrating that RPN1 specifically directs spike into the proteasomal degradation pathway. Given RPN1’s physical association with the ribosome–SEC61 translocon^22^, we next examined whether spike degradation involves ribosome-associated quality control (RQC). However, overexpression of LTN1, the central E3 ligase of the RQC pathway^32^, failed to deplete spike (Extended Data Fig. 6c), arguing against co-translational targeting and instead supporting a post-translational ERAD mechanism.

**Fig. 3.**
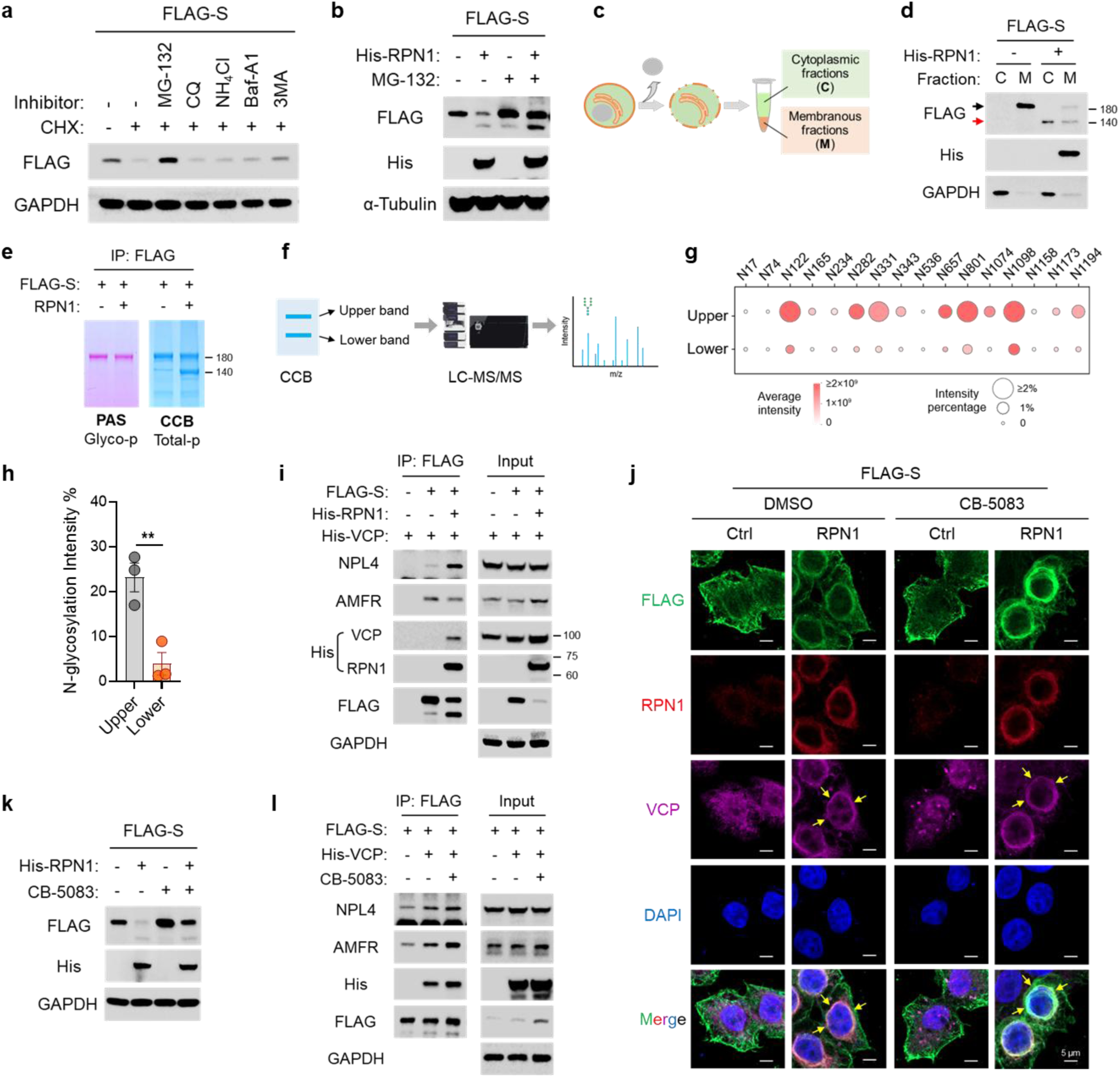
RPN1 induced the VCP/p97 complex-mediated ERAD of spike. **a**, Immunoblot analysis of S protein under proteasome or lysosomal inhibitor treatment. HEK293T cells overexpressing FLAG-S protein were treated with 20 μM MG-132, 50 μM chloroquine (CQ), 20 mM ammonium chloride (NH₄Cl), 50 μM bafilomycin A1 (Baf-A1), and 2 mM 3-methyladenine (3-MA) for 12 h, followed by 100 μg/mL cycloheximide (CHX) treatment for 2 h. **b**, Immunoblot analysis of S protein in cells co-expressing FLAG-S with His-RPN1. Cells were transfected for 24 h followed by 20 μM MG-132 treatment for 12 h. **c**, Schematic of cellular fractionation. Cells were lysed and separated into cytoplasmic (C) and membranous (M) fractions. **d**, Immunoblot analysis of S protein in C and M fractions from cells expressing FLAG-S with or without His-RPN1. The black and red arrows indicate the upper and lower bands of the S protein, respectively. **e**, Periodic acid-Schiff (PAS) staining and Coomassie brilliant blue (CBB) staining of total protein following immunoprecipitation of FLAG-S protein from HEK293T cells co-expressing FLAG-S with control or RPN1. PAS detects glycosylated proteins; CBB stains total protein. **f**, Schematic of LC-MS/MS analysis of S protein N-glycosylation. Upper and lower S protein bands from RPN1-overexpression cells were excised and analyzed. **g**, Site-specific N-glycosylation in S protein bands. Circle size represents the percentage of N-glycosylated peptide intensity and color denotes average N-glycosylation intensity. **h**, Quantification of total N-glycosylation intensity in the upper and lower S protein bands. Data are presented as mean ± SEM of biological replicates (n = 3), two-tailed unpaired *t*-test. (** *P* < 0.01) **i**, FLAG-S and control/His-RPN1 were co-transfected into the HEK293T overexpressing His-VCP. FLAG immunoprecipitation was performed, and co-precipitated proteins were probed with the indicated antibodies. **j**,**k**, FLAG-S and control/His-RPN1 were co-transfected into the HEK293T for 24 h, followed by treatment with DMSO or 1 μM CB-5083 for 12 h. (**j**) Immunofluorescence staining shows S (FLAG, green), RPN1 (red), VCP (magenta), and nuclei (DAPI, blue). (**k**) Immunoblotting shows changes in S protein levels. **l**, FLAG-S and control/His-VCP were co-transfected into the HEK293T for 24 h, followed by treatment with DMSO or 1 μM CB-5083 for 12 h. FLAG immunoprecipitation was performed, and co-precipitated proteins were probed with the indicated antibodies. GAPDH or α-Tubulin serves as a loading control (**a**, **b**, **d**, **i**, **k** and **l**).

The ERAD pathway is the best-understood organellar protein quality control system^33^. ERAD requires extraction of misfolded or regulated substrates from the ER lumen or membrane back into the cytosol for ubiquitination and proteasomal degradation^34^. Subcellular fractionation followed by immunoblotting revealed that RPN1 overexpression induced a redistribution of spike. While control cells displayed a single, full-length, membrane-localized spike band (∼180 kDa), RPN1-overexpressing cells exhibited two distinct bands, a weak ∼180 kDa band in the membrane fraction and a novel, lower-molecular-weight (LMW) band (∼130–140 kDa) both in the membrane and cytosolic fractions (Fig. 3c,d and Supplementary Table 1). Because spike’s theoretical molecular weight (<140 kDa) is increased to ∼180 kDa in mammalian cells by extensive N-linked glycosylation^35^, we hypothesized that the LMW band represents no-glycosylated spike following retrotranslocation. Periodic acid–Schiff (PAS) staining, which detects glycoproteins, confirmed that the ∼180 kDa band was strongly glycosylated, whereas the LMW band showed negligible PAS signal, while coomassie blue (CCB) staining verified equal protein loading (Fig. 3e). Mass spectrometry analysis of immunopurified LMW spike identified a dramatic reduction in N-glycosylation intensity (Fig. 3f-h). To validate this, we generated a N-glycosylation-deficient spike mutant (13N-Q) by mutating all 13 N-glycosylation sites in the S1 domain to glutamine (N→Q); its electrophoretic mobility matched the RPN1-induced LMW band (Extended Data Fig. 7). Together, these data support that RPN1 drives the retrotranslocation of spike from the ER into the cytosol—a prerequisite for its subsequent ubiquitination and proteasomal degradation.

The ERAD pathway operates through multiple parallel branches, each organized around through distinct membrane-resident E3 ubiquitin ligases^36^. Although individual branches differ in their specific protein components and substrate repertoires, the core mechanistic steps-substrate recognition, retrotranslocation, ubiquitination, extraction and proteasomal degradation-are broadly conserved across species^37^. To identify the key regulator for ERAD of spike, we performed immunoprecipitation coupled with mass spectrometry (IP-MS) using spike as bait. This analysis identified valosin-containing Protein (VCP, also known as p97), a highly conserved, essential AAA+ ATPase that governs multiple proteostatic pathways^38,39^, including ERAD, as a robust interactor (Supplementary Table 2). VCP functions within a canonical heterotrimeric complex consisting of its cofactors NPL4 and UFD1, and can collaborate functionally with the ER membrane–embedded E3 ubiquitin ligase AMFR^36^. Co-immunoprecipitation (Co-IP) assays validated direct physical interactions between Spike and each component of this machinery—NPL4, UFD1, and AMFR—and further revealed that RPN1 specifically strengthens the spike–VCP interaction (Fig. 3i and Extended Data Fig. 8a). Using LSCM, we observed that spike accumulates at the perinuclear ER and co-localizes with relocolized VCP and NPL4 upon co-overexpression of spike and RPN1; however, VCP relocolization was not observed upon overexpression of spike or RPN1 alone (Fig. 3j and Extended Data Fig. 8b,c).

Critically, pharmacological inhibition of VCP with CB-5083, a potent, selective, and cell-permeable ATPase inhibitor^40^, suppressed RPN1-dependent spike degradation and concurrently increased spike–VCP complex co-immunoprecipitation and co-localization, consistent with blockade of retrotranslocation and accumulation of stalled substrate–VCP complexes at the ER membrane (Fig. 3j,l and Extended Data Fig. 8c). Collectively, these results establish that RPN1 promotes VCP complex-dependent ERAD of spike protein.

### AMFR is the E3 ligase of SARS-CoV-2 spike protein

To establish AMFR as the E3 ubiquitin ligase responsible for spike ubiquitination, we performed gain- and loss-of-function experiments. Overexpression of wild-type AMFR significantly reduced spike protein levels, whereas an enzymatically inactive AMFR double mutant (C356G/H361A^41^) failed to induce this reduction (Fig. 4a). Consistent with impaired spike degradation, pseudotyped virus infectivity was markedly decreased upon AMFR overexpression—and this effect was partially reversed by co-expression of the catalytically deficient mutant (Fig. 4b). Conversely, CRISPR-mediated knockout of AMFR increased both endogenous spike abundance and SARS-CoV-2pp infectivity (Fig. 4c,d). Furthermore, AMFR overexpression enhanced total ubiquitination of spike, with a pronounced increase specifically in K48-linked polyubiquitin chains—canonical signals for proteasomal targeting—whereas both the AMFR mutant and AMFR knockout substantially diminished K48-linked ubiquitination (Fig. 4e,f). Collectively, these data demonstrate that AMFR functions as the E3 ligase for spike, mediating its VCP-dependent retrotranslocation and subsequent K48-linked ubiquitin-proteasome degradation. And RPN1 acts upstream to facilitate this process by strengthening the spike–VCP interaction.

**Fig. 4.**
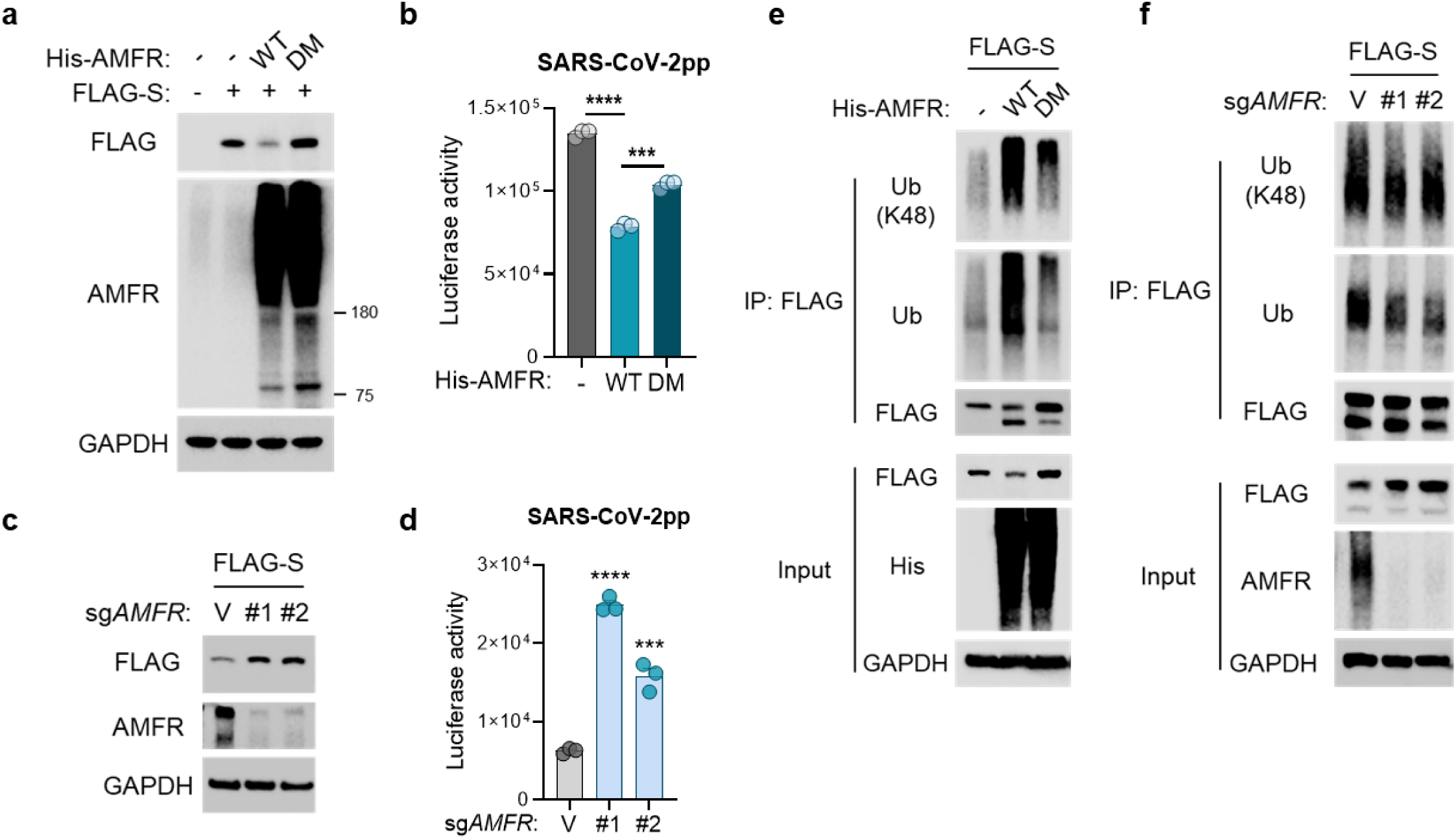
AMFR is the E3 ligase of spike. **a**, Immunoblot analysis of S protein in cells co-expressing FLAG-S with either His-AMFR (WT) or its double mutant (DM, C356G/H361A). **b**, Luciferase activity in Huh7 cells infected with SARS-CoV-2pp produced in control cells or in cells overexpressing AMFR (WT or DM). **c**, Immunoblot analysis of SARS-CoV-2 S protein in control or AMFR-knockout HEK293T cells. **d**, Luciferase activity in Huh7 cells infected with SARS-CoV-2pp produced in control or AMFR-knockout cells. **e**, FLAG-S and control/His-AMFR (WT or DM) were co-transfected into the HEK293T. FLAG immunoprecipitation was performed under denaturing conditions to preserve ubiquitin conjugates, followed by immunoblotting to detect K48-linked or total ubiquitination using the indicated antibodies. **f**, FLAG-S was transfected in control or AMFR-knockout HEK293T cells. FLAG immunoprecipitation was performed under denaturing conditions to preserve ubiquitin conjugates, followed by immunoblotting to detect K48-linked or total ubiquitination using the indicated antibodies. GAPDH serves as a loading control (**a**, **c**, **e** and **f**). Data are presented as mean ± SEM of biological replicates, two-tailed unpaired *t*-test of biological replicates (n = 3) (**b** and **d**). (*** *P* < 0.001; **** *P* < 0.0001)

### RPN1-mediated ERAD pathway shows conserved activity across highly pathogenic coronaviruses

SARS-CoV and MERS-CoV represent the other two highly pathogenic human coronaviruses that are phylogenetically distinct from SARS-CoV-2 and utilize different cellular receptors—SARS-CoV binds ACE2 with lower affinity than SARS-CoV-2, whereas MERS-CoV engages DPP4^42^. Co-expression of RPN1 with SARS-CoV or MERS-CoV spike proteins significantly reduced their levels in HEK293T cells (Fig. 5a), and concurrently enhanced physical association between each spike and endogenous VCP—as confirmed by co-immunoprecipitation (Extended Data Fig. 9a). Notably, while MERS-CoV spike robustly induces syncytium formation, SARS-CoV spike exhibits minimal syncytium-inducing capacity under identical experimental conditions^13,31,43,44^. Consistent with this functional distinction, RPN1 expression specifically suppressed MERS-CoV spike–induced syncytium formation (Fig. 5b,c, Extended Data Fig. 9b and Supplementary Videos 3-4). Furthermore, pseudotyped virus production assays demonstrated that RPN1 overexpression markedly decreased spike protein incorporation into both SARS-CoV pseudotyped particle (SARS-CoVpp) and MERS-CoV pseudotyped particle (MERS-CoVpp), leading to significantly reduced transduction efficiency in ACE2-expressing HeLa-ACE2 cells (for SARS-CoVpp) and Huh-7 cells (for MERS-CoVpp) (Fig. 5d-j). Collectively, these cross-coronavirus analyses demonstrate that RPN1’s capacity to target coronavirus spike proteins for VCP-dependent ERAD is not SARS-CoV-2–specific but constitutes a broadly conserved, receptor-agnostic host defense mechanism against multiple clinically significant coronaviruses.

**Fig. 5.**
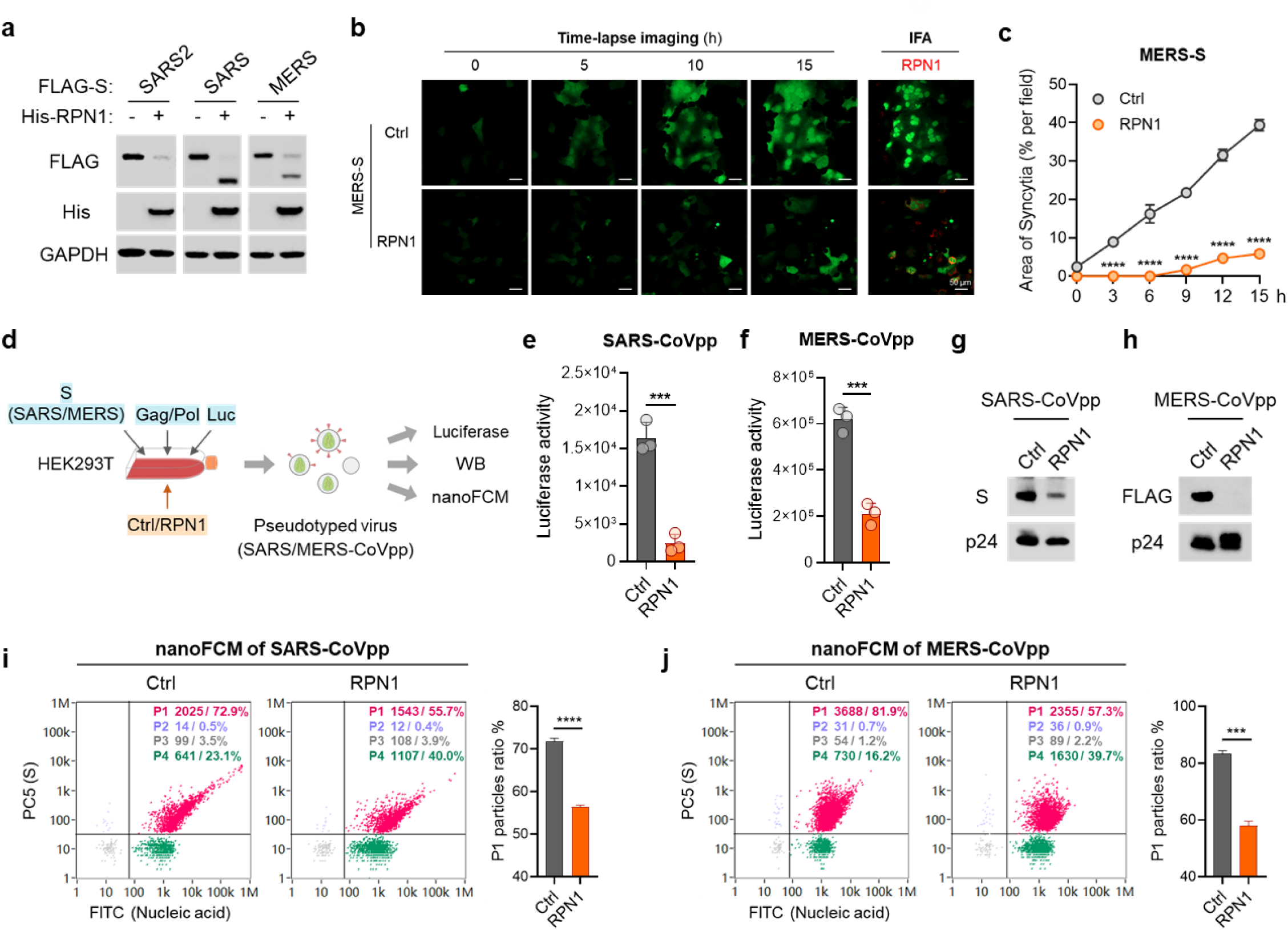
RPN1-mediated ERAD pathway shows conserved activity across highly pathogenic coronaviruses. **a**, Immunoblot analysis of S protein of SARS-CoV-2 (SARS2), SARS-CoV (SARS), and MERS-CoV (MERS) in HEK293T cells with or without RPN1 overexpression. GAPDH serves as a loading control. **b**,**c**, Huh7 cells were co-transfected with plasmids encoding S protein of MERS-CoV, eGFP, and control or RPN1. Live-cell time-lapse imaging was initiated 6 h post-transfection and continued for 15 h. See Supplementary Videos 3-4 for time-lapse movie. (**b**) Representative frames at 0, 5, 10, and 15 h into imaging show eGFP-labeled cell-cell fusion; after fixation at 15.5 h, immunofluorescence staining for RPN1 (red) is shown on the right. (**c**) Syncytial area per field was quantified every 3 h from time-lapse recordings. **d**, Schematic of the SARS-CoV pseudotyped particle (SARS-CoVpp) and MERS-CoV pseudotyped particle (MERS-CoVpp) production and analysis. **e**-**j**, Pseudotyped virus were generated in HEK293T cells by co-transfection of plasmids encoding SARS-CoV or MERS-CoV S, Gag/Pol, and Luc with or without of RPN1. Pseudotyped particles were purified for further analyzation. (**e** and **f**) Luciferase activity measured in Huh7 cells infected with SARS-CoVpp (**e**) or MERS-CoVpp (**f**) (n = 3). (**g** and **h**) Immunoblot analysis of S protein in SARS-CoVpp (**g**) and MERS-CoVpp (**h**); SARS-CoVpp was detected with an S antibody and MERS-CoVpp with a FLAG antibody; HIV Gag-p24 serves as a loading control. **i**,**j**, NanoFCM analysis of SARS-CoVpp (**i**) and MERS-CoVpp (**j**); Representative density plots show pseudotyped particle populations stained for S staining (PC5) and nucleic acid staining (FITC); Four gates (P1–P4) were defined; P1 (red) corresponds to RNA⁺S⁺ intact particles. Percentages of events in each gate are indicated. Right: quantitative comparison of P1 particle frequency between control and RPN1-overexpressing conditions. Data are presented as mean ± SEM, two-tailed unpaired *t*-test (**c**, **e**, **f**, **i** and **j**). (*** *P* < 0.001; **** *P* < 0.0001). All experiments were repeated at least three times independently.

### RPN1 is a promising therapeutic target for inhibiting SARS-CoV-2 infection

RPN1 is a type I transmembrane protein comprising an N-terminal cytoplasmic domain, a single transmembrane helix, and a C-terminal ER luminal domain^21^. To map the domain responsible for spike ERAD, we generated a series of RPN1 truncation constructs fused to an N-terminal ER signal peptide (to ensure luminal targeting) and assessed their effects on spike stability (Fig. 6a). We found that the construct containing the ER luminal domain (aa 24–438) robustly promoted spike degradation and exhibited strong co-immunoprecipitation with spike (Fig. 6b and Extended Data Fig. 10). Further systematic subdivision of the luminal domain into three fragments—aa 24–179 (domain 01), aa 180–307 (domain 02), and aa 308–438 (domain 03)—revealed that domain 02 alone was both necessary and sufficient to drive spike ERAD, as evidenced by its potent reduction of spike protein levels relative to controls and other fragments (Fig. 6c). Consistent with this, overexpression of RPN1 domain 02 (RPN1^02^) in HeLa-ACE2 cells significantly suppressed spike-mediated syncytium formation and inhibited SARS-CoV-2 infection (Fig. 6d,e). These observations highlight the RPN1 luminal domain, particularly domain 02, as a compact and functionally independent antiviral module with promising translational potential. Targeting or delivering this minimal functional region may represent a novel and feasible strategy for developing host-directed, ERAD-based antiviral interventions.

**Fig. 6.**
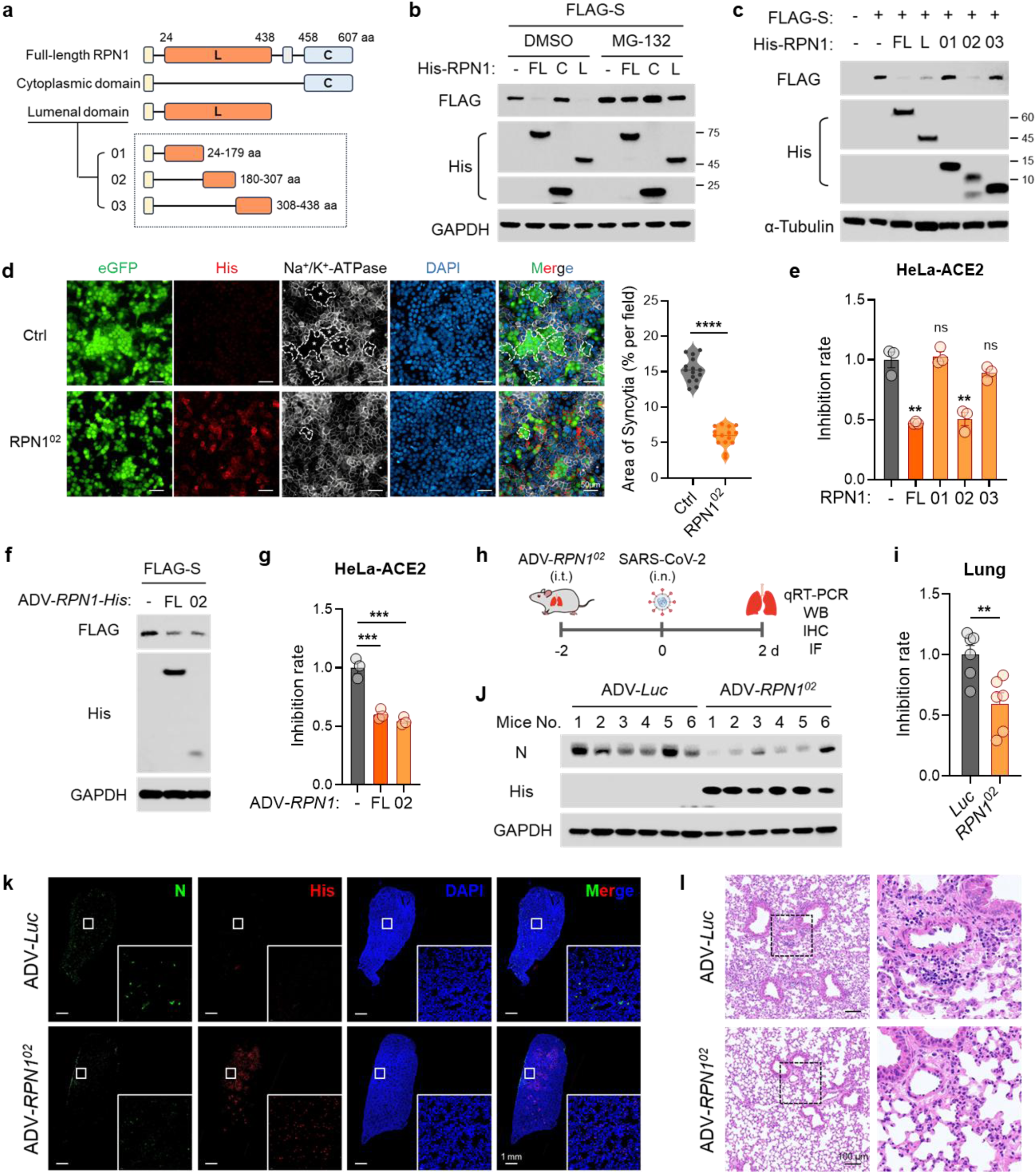
RPN1 is a promising therapeutic target for inhibiting SARS-CoV-2 infection. **a**, Schematic representation of the RPN1 domain architecture and corresponding truncation constructs. L, luminal domain; C, cytoplasmic domain. Truncations 01-03 encompass residues 24-179, 180-307, and 308-438, respectively. **b**, Immunoblot analysis of S protein in cells co-expressing FLAG-S with His-RPN1 full-length (FL), cytoplasmic domain (C), or luminal domain (L). Cells were transfected for 24 h followed by DMSO or 20 μM MG-132 treatment for 12 h. **c**, Immunoblot analysis of S protein in cells co-expressing FLAG-S with His-RPN1 (FL, L, or truncations 01-03). **d**, Immunofluorescence analysis of S-mediated syncytium formation in HeLa-ACE2 cells transfected with control or RPN1^02^. eGFP (green) was used to visualize S-mediated membrane fusion. RPN1 (red), Na⁺/K⁺-ATPase (membrane marker, gray), and nuclei (DAPI, blue) are shown. Dashed outlines with asterisks indicate representative syncytia. Quantification of total syncytial area per field is presented on the right. **e**, Quantification of viral RNA levels in HeLa-ACE2 cells overexpressing RPN1 FL or truncations following SARS-CoV-2 infection (MOI=0.01) for 24 h (n = 3). **f**, Immunoblot analysis of S protein expression in HeLa-ACE2 cells transduced with ADV-*RPN1^FL^-His* or ADV-*RPN1*^02^*-His* (MOI=40) for 24 h. **g**, HeLa-ACE2 cells were transduced with ADV-*RPN1 FL* or ADV-*RPN1*^02^ (MOI=40) for 24 h prior to infection with SARS-CoV-2 (MOI=0.01) for an additional 24 h. Viral RNA levels in culture supernatants, quantified by qRT-PCR and normalized to the average value of control group (set to 1), are shown as fold change relative to control (n = 3). **h**, Schematic of adenoviral vector delivery and SARS-CoV-2 infection in mice. **i**-**l**, BALB/c mice (n=6 per group) were intratracheally transduced with ADV*-Luc* or ADV*-RPN1*^02^ (2×10^9^ PFU) for 2 d, then infected intranasally with SARS-CoV-2 Beta variant (1×10^4^ PFU). Lungs were harvested at 2 dpi for downstream analysis. (**i**) Viral RNA levels, quantified by qRT-PCR, are presented relative to the ADV-*Luc* control group (set to 1) to calculate the percent inhibition conferred by ADV-*RPN1*^02^. (**j**) Immunoblot analysis of viral N protein and RPN1^02^ expression in lung lysates. (**k**) Immunofluorescence staining of lung sections; viral N protein (green), Rpn1 (His, red), and nuclei (DAPI, blue) are shown; boxed regions are enlarged in the bottom-right insets. (**l**) H&E staining of lung tissues. Insets display higher-magnification images of the boxed regions. GAPDH or α-Tubulin serves as a loading control (**b**, **c**, **f** and **j**). Data are presented as mean ± SEM, two-tailed unpaired *t*-test (**d**, **e**, **g** and **i**). (ns, not significant; ** *P* < 0.01; *** *P* < 0.001; **** *P* < 0.0001). Data represent at least three independent experiments.

### Therapeutic efficacy of recombinant RPN1 against SARS-CoV-2

To evaluate the therapeutic potential of RPN1 *in vivo*, we first packaged the full-length or domain 02 coding sequence into a conditionally replicative adenoviral vector (ADV-*RPN1* or ADV-*RPN1*^02^). Both constructs were efficiently expressed in HeLa-ACE2 cells and significantly suppressed SARS-CoV-2 infection—consistent with their established roles in regulating spike ERAD (Fig. 6f,g). Finally, to validate the therapeutic effect of RPN1, ADV-RPN1^02^ was administrated to the lungs of mice via intratracheal (i.t.) instillation 2 days prior to SARS-CoV-2 challenge (Fig. 6h). Recombinant adenovirus expressing Luciferase (ADV-Luc) was used as control. Compared with ADV-Luc, ADV-RPN1^02^ administration readily decreased pulmonary viral RNA load and N protein levels (Fig. 6i-k), as well as attenuated SARS-CoV-2–induced infiltration of lung immune cells (Fig. 6l). Collectively, these results demonstrate that RPN1^02^ is a functionally autonomous, ER luminal domain that drives spike ERAD and exerts potent, mechanism-based antiviral activity both *in vitro* and *in vivo*.

## Discussion

In this study, we identify RPN1 as a previously uncharacterized, spike-selective ERAD factor. Following co-translational translocation into the ER lumen, the coronavirus spike protein is specifically recognized by the ER luminal domain of RPN1, triggering AMFR-mediated K48-linked ubiquitination. This marks spike for VCP/p97-dependent retrotranslocation and proteasomal degradation. By depleting functional spike from the ER pool, this RPN1-initiated ERAD pathway markedly reduces spike incorporation into progeny virions and suppresses spike-driven syncytium formation in infected cells—thereby attenuating both cell-to-cell viral spread and spike-associated cytopathic effects.

ERAD, a critical cellular protein quality control machinery, is tightly involved in the life cycles of multiple viruses (e.g. flaviviruses, HIV, and coronaviruses), and is frequently co-opted by these pathogens to support viral entry, replication, and evasion of host immune surveillance^45,46^. In coronaviruses, the non-structural protein 6 (nsp6) hijack ERAD to facilitate lipolysis for the growth of double-membrane vesicles (DMVs)^47^. However, despite ERAD’s well-established role as a host defense mechanism for eliminating misfold or aberrant proteins, evidence remains limited regarding whether the host exploits ERAD to selectively target and degrade coronavirus-encoded proteins as a means of restricting infection. In this study, we identify RPN1 as an ERAD factor that restricts coronavirus infectivity—and as a potential therapeutic target for combating highly pathogenic coronaviruses. Prior to this work, the experimentally validated role for RPN1 in viral replication was its requirement for the N-glycosylation of the Dengue virus (DENV) non-structural protein 1 (NS1)^48^, a modification critically required for the infectivity of DENV and some other viruses such as SARS-CoV-2^28,45^. In contrast, our study identifies RPN1 as a previously unrecognized antiviral factor that functions directly within the ERAD pathway to selectively restrict coronavirus infection, thereby revealing a novel axis of ERAD–virus crosstalk and establishing a mechanistic basis for the development of host-directed antivirals targeting ERAD components.

RPN1’s strategic localization at the ribosome–SEC61 translocon interface uniquely positions it to monitor nascent polypeptides as they enter the ER. As a core subunit of OSTs, RPN1 plays an essential role in the co-translational N-glycosylation and membrane integration of newly synthesized polypeptides^22^. Its proximity to the entry point of the secretory pathway underlies its non-redundant physiological function. We attempted to generate global Rpn1 knockout mice, however, this intervention results in early embryonic lethality (data not shown). This lethality precludes conventional loss-of function studies in adult animals, necessitating conditional, cell-type–specific deletion for *in vivo* functional analysis.

Although our study focuses on spike protein, we cannot exclude the possibility that RPN1’s ERAD activity extends to other misfolded or N-glycosylated membrane proteins —particularly those harboring exposed luminal domains or transmembrane segments susceptible to retrotranslocation. Whether the spike-targeting ERAD activity of RPN1 intersects with or affects its canonical function in OST-mediated N-glycosylation remains to be fully elucidated. Nevertheless, RPN1 represents a promising host-directed therapeutic target, owing to its selective antiviral action toward spike. Importantly, our results identify the RPN1 luminal domain 02 (residues 180–307) as a minimal, autonomous antiviral module that retains full activity without disrupting global cellular glycosylation, offering a translatable blueprint for peptide- or gene-based antiviral development.

Overall, the identification of RPN1 as a dual-function protein—serving both canonical glycosylation and antiviral ERAD roles—exemplifies how cellular machinery can be evolutionarily repurposed for pathogen recognition. This finding suggests that systematic investigation of other ER-resident enzymes for antiviral activities could reveal additional layers of innate immune defense and provide new targets for therapeutic intervention.

## Methods

### Ethics and biosafety

All animal studies were performed in strict accordance with the guidelines set by the Chinese Regulations of Laboratory Animals and Laboratory Animal-Requirements of Environment and Housing Facilities. All animal procedures were reviewed and approved by the Animal Experiment Committee of Laboratory Animal Center, Academy of Military Medical Sciences (AMMS) (approval number: IACUC-IME-2022-051) or the Guangzhou Customs District Technology Center (approval number: IQTC2025004-01).

### Cells

Cell lines including HEK293T (embryonic kidney, ATCC CRL-11268), HeLa-ACE2, Caco2 (colorectal adenocarcinoma, BNCC350769), HeLa (cervical carcinoma, ATCC CCL-2), Huh7 (hepatocellular carcinoma, Procell CL-0120), Vero (African green monkey kidney, ATCC CCL-81) were cultured in Dulbecco’s Modified Eagle’s Medium (DMEM, Gibco 11995065) supplemented with 10% fetal bovine serum (FBS, Gibco A5670701) and 1% penicillin-streptomycin (P/S, Gibco 15140122) at 37°C in a humidified incubator with 5% CO_2_. All cell lines were routinely tested and confirmed negative for mycoplasma contamination before use.

### Viruses

The SARS-CoV-2 strain BetaCov/human/CHN/Beijing_IME-BJ05/2020 (IME-BJ05, accession no. GWHACAX01000000) was used as the WT virus in this study. SARS-CoV-2 WT and the Beta (GDPCC, CSTR: 16698.06. NPRC2.062100001), Delta (CCPM-B-V-049-2105-8), and JN.1(GB0009319, accession: C_AA160427.1) variants were preserved in Biosafety Level 3 facilities of Academy of Military Medical Sciences (AMMS). Beta (B.1.315, strain: No.2021XG-888) for *Rpn1*-cKO mice was preserved in Biosafety Level 3 facilities of Guangzhou Customs District Technology Center. All SARS-CoV-2 viral stocks were propagated in Vero cells and titrated by plaque-forming assay (for WT, Beta and Delta) or focus-forming assay (FFA, for JN.1) and confirmed by next-generation sequencing. All experiments involving infectious SARS-CoV-2 were performed under Biosafety Level 3 facilities in AMMS or the Guangzhou Customs District Technology Center.

### Antibodies

The following primary antibodies were used in this study: mouse monoclonal anti-SARS-CoV-2 nucleocapsid antibody (Sino Biological, 40143-MM08), mouse monoclonal anti-SARS-CoV/SARS-CoV-2 spike antibody (Sino Biological, 40150-D001), rabbit monoclonal anti-MERS-CoV spike antibody (Sino Biological, 40069-R723), rabbit monoclonal anti-SARS-CoV-2 spike antibody (Sino Biological, 40592-R0004), rabbit polyclonal anti-HIV-1 Gag-p24 antibody (Sino Biological, 11695-RP02), rabbit polyclonal anti-RPN1 antibody (ABclonal, A6726), rabbit polyclonal anti-Rpn1 antibody (ABclonal, A12497), mouse monoclonal anti-His-tag antibody (ABclonal, AE003), rabbit monoclonal anti-VCP antibody (ABclonal, A1402), rabbit monoclonal anti-Na⁺/K⁺-ATPase antibody (ABclonal, A11683), rabbit monoclonal anti-ubiquitin antibody (ABclonal, A19686), rabbit monoclonal anti-K48-linkage-specific ubiquitin antibody (ABclonal, A3606), rabbit monoclonal anti-His-tag antibody (Cell Signaling Technology, 12698), rabbit monoclonal anti-β-actin antibody (Cell Signaling Technology, 4970), rabbit monoclonal anti-GAPDH antibody (Cell Signaling Technology, 2118), rabbit monoclonal anti-HA-tag antibody (Cell Signaling Technology, 3724), rabbit polyclonal anti-NPL4 antibody (Cell Signaling Technology, 13489), mouse monoclonal anti-FLAG antibody (Sigma-Aldrich, F3165), mouse monoclonal anti-α-tubulin antibody (Sigma-Aldrich, T5168) and rabbit polyclonal anti-AMFR antibody (Proteintech, 16675-1-AP).

The following secondary antibodies were used: CoraLite 488-conjugated donkey anti-mouse IgG (Proteintech, SA00013-5), CoraLite 594-conjugated donkey anti-rabbit IgG (Proteintech, SA00013-7), Multi-rAb CoraLite Plus 647-conjugated goat anti-rabbit IgG (Proteintech, RGAR005), horseradish peroxidase (HRP)-conjugated goat anti-rabbit IgG (Jackson ImmunoResearch, 111-035-003) and HRP-conjugated goat anti-mouse IgG (Jackson ImmunoResearch, 115-035-003).

### Rpn1^fl/fl^, Sftpc^CreERT2^ *mice*

*Rpn1^fl/fl^, S*ftpc*^CreERT2^*mice were generated by Cyagen Biosciences (Suzhou, China). In brief, CRISPR/Cas9-based editing was used to insert loxP sites around exon 3 of the Rpn1 gene. The gRNA to mouse Rpn1 gene, the donor vector containing loxP sites, and Cas9 mRNA were co-injected into fertilized mouse eggs to generate targeted conditional knockout offspring. F0 founder animals were identified by PCR followed by sequence analysis, which were bred to wildtype mice to test germline transmission and F1 animal generation. Breed a F1 targeted mouse with a type II alveolar epithelial cells-specific Sftpc-CreERT2 delete mouse to generate FN. *Rpn1^fl/fl^* mice were crossed to *Rpn1^fl/fl^, S*ftpc*^CreERT2^*mice. Experiments included roughly equal numbers of male and female animals and were pooled from multiple litters. Animals of both genotypes were cohoused.

### Animal models

Specific pathogen-free 8 weeks old male BALB/c mice were purchased from Beijing Vital River Laboratory Animal Technology Co., Ltd (Beijing, China). *Rpn1^fl/fl^, S*ftpc*^CreERT2^*mice were generated from the C57BL/6J strain and provided by the Cyagen Biosciences (Suzhou, China). *Rpn1^fl/fl^, S*ftpc*^CreERT2^*mice were administered tamoxifen (MedChemExpress, HY-13757A) dissolved in corn oil (MedChemExpress, HY-Y1888) by intraperitoneal injection at 100 mg/kg once daily for 5 consecutive days. Tamoxifen injection was administered 7 weeks before SARS-CoV-2 infection. Animals were kept under specific-pathogen-free conditions, with controlled temperature (21°C–23°C), humidity (30–60%) and light cycle (12 h light/dark).

### Virus infection of mice

*Rpn1^fl/fl^* and *Rpn1^fl/fl^, S*ftpc*^CreERT2^*male mice (7 months old) were infected intranasally with SARS-CoV-2 Beta variant (B.1.315, 1×10^4^ FFU per mouse) in a total volume of 20 μL PBS. Mice were sacrificed at 3 days post-infection. Serum samples were collected for viral RNA quantification, and lung tissues were harvested for FFA, RNA extraction, immunoblotting, immunofluorescence staining, and histopathological examination. Samples for RNA analysis were preserved in TRIzol reagent (Invitrogen, 15596026); tissues for immunoblotting were lysed in RIPA (no SDS) buffer; tissues for histology were fixed in 4% paraformaldehyde (PFA) and processed for hematoxylin-eosin (H&E) staining.

Syncytial pneumocytes in lung sections of SARS-CoV-2 infected mice were quantified on H&E staining using ImageJ with an unbiased sampling. Random non-overlapping regions of interest (ROIs) were generated across the lung section using a custom ImageJ macro, a minimum spacing of one high-magnification field between adjacent ROIs. For each mouse lung, at least 20 high-magnification ROIs were randomly selected and analyzed according to American Thoracic Society (ATS) guidelines. ROIs containing large airways or vessels or consisting of <50% lung parenchyma were excluded prior to analysis. The selected ROIs were subsequently used for quantitative syncytia assessment. Syncytia were defined as multinucleated pneumocytic aggregates exhibiting (i) cytoplasmic continuity without discernible cell boundaries and (ii) nuclear clustering within an expanded cytoplasmic domain. Syncytia area (%) was calculated as the percentage of ROI area.

### Plasmid transfection and infection

HeLa-ACE2 cells were seeded in 12-well plates or T75 flasks and transfected the following day with pcDNA3.1-RPN1-His plasmid or control plasmid using Lipofectamine^TM^ 3000 (Thermo Fisher Scientific, L3000015) according to the manufacturer’s instruction. At 24 h post-transfection, cells were infected with SARS-CoV-2 at a multiplicity of infection (MOI) of 0.01. Culture supernatants were collected at 24 h or 48 h post-infection for viral RNA quantification by quantitative PCR (qRT-PCR), and cell lysates were harvested at 48 h for immunoblot analysis using antibodies against SARS-CoV-2 nucleocapsid (mouse monoclonal, Sino Biological, 40143-MM08, 1:1000) and His (rabbit monoclonal, Cell Signaling Technology, 12698, 1:1000). Immunofluorescence staining was performed at 48 h as described in the Immunofluorescence section.

### RNA Interference and infection

Vero and Caco2 cells were seeded in 12-well plates and transfected with siRNAs at a final concentration of 100 nM using Lipofectamine^TM^ RNAiMAX (Thermo Fisher Scientific, 13778150). Human *RPN1* (stB0001906B and stB0001906C), African green monkey *RPN1* (siG2111231130329569 and siG2111231130330661) and NC (siN0000001) siRNAs were purchased from RiboBio. At 48 h post-transfection, cells were infected with SARS-CoV-2 at a MOI of 0.01. Culture supernatants were collected for viral RNA quantification by qRT-PCR, and cell lysates were harvested for immunoblot analysis using antibodies against SARS-CoV-2 nucleocapsid (mouse monoclonal, Sino Biological, 40143-MM08, 1:1000) and RPN1 (rabbit polyclonal, ABclonal, A6726, 1:1000). The siRNA sequences are as follows: Human *RPN1* siRNA: #1: GCTACAACCTCCCAAGCTA; #2: CTCGAAATGTGGAGAGCTA. Chlorocebus sabaeus *RPN1* siRNA: #1: AGCTCAAGAAAGACACGTA; #2: CTGGACTTCTCCATCACTA.

### Western blot

Cells were harvested in lysis buffer supplemented with 1 mM PMSF and protease inhibitor cocktail (Roche, 04693132001). Samples were separated by Precast-Glgel 4-20% Tris-Glycine PAGE (Sangon Biotech, C661105-0001) and transferred to 0.45 µm PVDF membranes (Millipore, IPFL00010). Membranes were blocked with 5% non-fat milk (Sangon Biotech, A600669) for 1 h at room temperature and incubated with primary antibodies overnight at 4°C. Then, membranes were incubated with HRP-conjugated goat anti-rabbit or anti-mouse IgG secondary antibodies (Jackson ImmunoResearch, 111-035-003 or 115-035-003) for 1 h at room temperature. Protein bands were detected using SuperSignal™ West Pico PLUS Chemiluminescent Substrate (Thermo Fisher Scientific, 34580) and imaged with a ChemiDoc Touch Imaging System (Bio-Rad).

### Real-time quantitative PCR

Serum and lung tissues were subjected to RNA extraction using TRIzol reagent (Invitrogen, 15596026) according to the manufacturer’s instructions. For cell culture supernatants, RNA was isolated using the qEx-DNA/RNA Kit (TIANLONG, T335) following the recommended protocol. Viral genomic RNA was quantified by one-step quantitative reverse transcription PCR (RT-qRT-PCR) using the One Step PrimeScript RT-PCR Kit (Takara, RR064A) on a LightCycler^®^ 480 Real-Time PCR System (Roche). Absolute RNA copy numbers were calculated based on a standard curve generated from serial dilutions of synthetic RNA transcripts. The primer sets were as follows:

SARS-CoV-2 gRNA-F: ACAGGTACGTTAATAGTTAATAGCGT;

SARS-CoV-2 gRNA-R: ATATTGCAGCAGTACGCACACA;

SARS-CoV-2 gRNA-probe: FAM-ACACTAGCCATCCTTACTGCGCTTCG-BHQ1;

optimized S-F: TGTACTTCGCCAGCACCGAA;

optimized S-R: GTTCTTGTGGTAGTACACGCC;

WPRE-F: CCTTTCCGGGACTTTCGCTTT;

WPRE-R: GCAGAATCCAGGTGGCAACA.

### Immunofluorescence

HeLa cells were seeded onto glass slides to reach 30-40% confluence the following day and co-transfected with pcDNA3.1-3×FLAG-S-tStrep and pcDNA3.1-RPN1-His or control plasmids. At 24 h post-transfection, cells were fixed with 4% PFA for 10 min at room temperature and permeabilized with 0.2% Triton X-100 (Solarbio, T8200) for 10 min at 4°C. After blocking in 5% BSA for 1 h at room temperature, cells were incubated with primary antibodies against FLAG tag (mouse monoclonal, Sigma, F3165, 1:400) and VCP (rabbit monoclonal, ABclonal, P55072, 1:100) overnight at 4°C. Following washing, cells were incubated with CoraLite 488-conjugated Donkey Anti-Mouse (Proteintech, SA00013-5) and CoraLite 594-conjugated Donkey Anti-Rabbit (Proteintech, SA00013-7, 1:200) secondary antibodies for 1 h at room temperature. RPN1 antibody (rabbit polyclonal, ABclonal, A6726) was conjugated to Alexa Fluor 647 using an Alexa Fluor^®^ 647 Conjugation Kit (Abcam, ab269823) according to the manufacturer’s instructions and incubated with cells overnight at 4°C. Coverslips were mounted using a DAPI-containing mounting medium (Abcam, ab104139). Fluorescence images were acquired using a Zeiss LSM980 confocal laser scanning microscope equipped with a 40× oil-immersion objective.

For staining following viral infection, cells were fixed with 4% PFA and permeabilized with 0.2% Triton X-100 (Solarbio, T8200). After blocking with 5% BSA, cells were incubated with primary antibodies against SARS-CoV-2 nucleocapsid (mouse monoclonal, Sino Biological, 40143-MM08, 1:400) and RPN1 (rabbit polyclonal, ABclonal, A6726, 1:100) overnight at 4°C. Cells were then incubated with CoraLite Plus 647-Goat Anti-Rabbit (Proteintech, RGAR005) and CoraLite 594-conjugated donkey anti-mouse (Proteintech, SA00013-7, 1:200) secondary antibodies for 1 h at room temperature. Images were acquired using an Operetta CLS high-content screening system (PerkinElmer) equipped with a 20× objective.

### Cell-cell fusion assay

HeLa-ACE2 or Huh7 cells were seeded in 12-well plates to reach 60-70% confluence the following day. Cells were co-transfected with pcDNA3.1-3×FLAG-S-tStrep or pcDNA3.1-3×FLAG-MERS-CoV S and pcDNA3.1-T2A-eGFP plasmids in the presence or absence of pcDNA3.1-RPN1-His or pcDNA3.1-RPN1 02-His plasmid. The green fluorescence was used to monitor membrane fusion and syncytium formation. At 16-30 h post-transfection, cells were fixed with 4% PFA and permeabilized with 0.2% Triton X-100 (Solarbio, T8200). After blocking with 5% BSA, cells were stained with antibodies against Na⁺/K⁺-ATPase (ABclonal, A11683) as a membrane marker and RPN1 (ABclonal, A6726) or His-tag (ABclonal, AE003) antibody. Nuclei were stained with DAPI (Sigma, S7113).

For viral infection-induced cell-cell fusion assays, HeLa-ACE2 cells were seeded in 12-well plates to reach 50% confluence on the following day. Cells were co-transfected with pcDNA3.1-RPN1-His or pcDNA3.1-T2A-luciferase (control plasmid) and pcDNA3.1-T2A-eGFP plasmids. The green fluorescence was used to monitor membrane fusion and syncytium formation. At 24 h post-transfection, cells were infected with sars-Cov-2 Delta variant for 48 h. Subsequently, cells were fixed with 4% PFA and permeabilized with 0.2% Triton X-100 (Solarbio, T8200). After blocking with 5% BSA, cells were stained with antibodies against S (Sino Biological, 40592-R0004) and His-tag (ABclonal, AE003) antibody. Nuclei were stained with DAPI (Sigma, S7113).

Images were acquired using an Operetta CLS high-content screening system (PerkinElmer) equipped with a 20× objective. Image analysis was performed using Harmony software (v4.9, PerkinElmer). Images were first subjected to flat-field correction. For cells expressing green fluorescence were segmented using the ‘Find Cells’ analysis module based on the 488-nm channel. Nuclei were identified within the positive green signal using the ‘Find Nuclei’ module. The thresholds for image segmentation were adjusted according to the signal-to-background ratio. Splitting coefficient was set to avoid splitting of overlapping nuclei. The cytoplasm area was defined using the ‘Find Cytoplasm’ module based on green fluorescence. Multinucleated cells were identified using Method E. For each cell, the number of nuclei and cell area were calculated. For MERS-CoV S-mediated membrane fusion, cells expressing green fluorescence were segmented using the ‘Find Image Region’ analysis module based on the 488-nm channel. Nuclei were identified within ‘Image Region’ using the ‘Find Nuclei’ module. Image regions containing more than four nuclei and exhibiting cytoplasmic green fluorescence were considered as fused cells. Syncytium formation was quantified as the percentage of total field area occupied by fused cells^1^.

For live-cell imaging, cells were seeded in 12-well glass-bottom plates (Cellvis, P12-1.5H-N). At 6 h post-transfection, time-lapse imaging was initiated using a Zeiss LSM980 confocal laser scanning microscope equipped with a 20× objective under 37°C and 5% CO_2_. Three fields per well were imaged every 20 min for 36 h (SARS-CoV-2) or 15 h (MERS-CoV) to monitor S protein-mediated membrane fusion, using excitation at 460-490 nm and emission at 500-550 nm. Image analysis was performed using Fiji. Cells exhibiting uniform green fluorescence throughout the cytoplasm were considered as fused cells. Syncytium formation was quantified as the percentage of total field area occupied by fused cells.

### SARS-CoV-2 plaque-forming assay

Viral supernatants were serially diluted in PBS and inoculated (100 μL per well) onto Vero cells seeded in 6-well plates. After adsorption for 1 h at 37°C with gentle rocking every 15 min, the inoculum was removed and cells were overlaid with 1% agarose containing 2% FBS. At 4 days post-infection, cells were fixed with 4% PFA and stained with 0.1% crystal violet for plaque visualization and enumeration.

### Focus forming assay (FFA)

Vero cells were seeded in 96-well plates one day prior to infection. Supernatants were serially diluted in PBS and used to inoculate cells at 37°C for 1 h. After removing the inoculum, plates were overlaid 125 μL pre-warmed 1.6% carboxymethylcellulose containing 2% FBS. After further incubation for 3 days, cells were fixed with 4% PFA and permeabilized with 0.2% Triton X-100 (Solarbio, T8200). Cells were then incubated with primary antibodies against SARS-CoV-2 nucleocapsid (mouse monoclonal, Sino Biological, 40143-MM08, 1:400), followed by an HRP-labeled goat anti-mouse secondary antibody (Jackson ImmunoResearch, 115-035-003, 1:5000). Viral foci were visualized by TrueBlue Peroxidase Substrate (SeraCare, 5510-0030), and counted with an ELISPOT reader (Cellular Technology Ltd. Cleveland, OH). Viral titers were calculated as focus-forming units (FFU) per ml.

### SARS-CoV-2 purification

To pellet authentic virions, supernatants were first clarified by centrifugation at 1,000 rpm for 3 min to remove cellular debris and subsequently inactivated with 4% PFA for 48 h at 4°C. The fixed virions were pelleted through a 20% sucrose cushion by ultracentrifugation (Beckman) at 100,000 g for 2 h at 4°C and resuspended in HEPES-saline buffer (10mM HEPES, 150mM NaCl) overnight at 4°C. Samples were stored at 4°C until further processing.

### Cryo-electron microscopy (cryo-EM)

4 μL of virus sample was rapidly applied to a glow-discharged Quantifoil (Au R1.2/1.3, 300 mesh carbon grid). On a Vitrobot Mark IV (FEI), the grid was blotted with filter paper for 3 s (8°C, 100% humidity) to remove excess sample. After waiting for 15 s, the grid was immersed in liquid ethane (-180°C) cooled by liquid nitrogen. Cryo-EM images were acquired at 78,000× magnification using an FEI Falcon III Direct Electron Detector on a Tecnai Arctica microscope.

### Adenoviral vector transduction in cell

Conditionally replicative adenoviral vectors expressing luciferase (ADV*-Luc*), full-length RPN1 (ADV*-RPN1^FL^*), and RPN1 180-307 aa domain (ADV*-RPN1*^02^) with a His tag were obtained from OBiO Technology (Shanghai, China). For plasmid expression assays, HeLa-ACE2 cells were first transfected with plasmids for 24 h, followed by transduction with ADV-*Luc*, ADV-*RPN1^FL^* or ADV-*RPN1*^02^ at MOI 40 for an additional 24 h. Protein expression was confirmed by immunoblotting. For viral infection experiments, HeLa-ACE2 cells were transduced with ADV-*Luc*, ADV-*RPN1^FL^*or ADV-*RPN1*^02^ at MOI 40 for 24 h and subsequently infected with SARS-CoV-2 at MOI 0.01 for 24 h. Cell culture supernatants were harvested for viral RNA quantification.

### Adenoviral vector transduction and infection in mice

8-week-old male BALB/c mice (n=6 per group) were anesthetized with sodium pentobarbital and transduced intratracheally with 2×10^9^ PFU of ADV-*Luc* or ADV-*RPN1*^02^ in a total volume of 40 μL PBS. Mice were intranasally infected with SARS-CoV-2 Beta variant (1×10^4^ PFU) 2 days post-transduction. All animals were sacrificed at 2 days post infection. Lung tissues were harvested for RNA extraction, immunoblot analysis, immunofluorescence staining, and histopathological examination. Samples for RNA analysis were preserved in TRIzol reagent (Invitrogen, 15596026), whereas tissues for histology were fixed in 4% PFA and processed for H&E staining.

### CRISPR-Cas9-mediated genome editing

Small guide RNAs (sgRNAs) were cloned into the lenti-CRISPRv2 vector and co-transfected with psPAX2 packaging plasmid (Addgene, 12260) and pMD2.G envelope plasmid (Addgene, 12259) into HEK293T cells. Viral supernatants were collected at 72 h post-transfection and filtered through a 0.45 μm membrane. Target cells were transduced with lentivirus in the presence of 8 μg/mL polybrene for 24 h and selected with puromycin to establish stable cell populations. Genome editing efficiency was confirmed by immunoblotting. The sgRNAs targeting AMFR (Genbank NM_001144.6) were designed as follows:

#1-F: CACCGCGTTAGCTGGTCCGGCTCGC;

#1-R: AAACGCGAGCCGGACCAGCTAACGC;

#2-F: CACCGTGGCTGCTCATCGGCGTGGT;

#2-R: AAACACCACGCCGATGAGCAGCCAC.

### Pseudotyped virus preparation, purification and infection

HEK293T cells were transfected with lentiviral reporter plasmid expressing luciferase (pLenti-V5-luciferase), psPAX2, and S expression plasmid (pcDNA3.1-S Δ19aa-3×FLAG) at a ratio of 6:3:1. Based on lentiviral system, another plasmid including control vector, pcDNA3.1-RPN1-His, or pCMV-AMFR-6×His-Neo (WT or DM) plasmid was co-transfected with S plasmid at a ratio of 2:1. Supernatant containing pseudotyped virus was harvested at 48 h post-transfection, filtered through a 0.45 μm membrane, and used immediately for infection. Pseudotyped virus production efficiency was assessed by quantifying lentiviral genome copies targeting WPRE sequence in the lentiviral reporter plasmid.

To pellet pseudotyped particles, viral supernatants were ultracentrifuged through a 20% sucrose cushion at 100,000×g for 2 h at 4°C using a Beckman SW32 rotor. Virus pellets were resuspended in PBS buffer for analysis of S protein incorporation and particle integrity through immunoblotting and nanoscale flow cytometry.

For infection assays, Huh7 cells were seeded in 24-well plates one day prior to infection and inoculated with equal amounts of pseudotyped virus. After 6-12 h incubation, medium was replaced with fresh culture medium. About 72 h post-infection, cells were lysed using Luciferase Assay System (Promega, E1501). Luminescence was measured using a GloMax^®^ Navigator Microplate Luminometer (Promega).

### Nanoscale flow cytometry

Sample was resuspended in PBS to a final volume of 50 μL. Alexa Fluor 647-conjugated anti-SARS-CoV/SARS-CoV-2 S (mouse monoclonal, Sino Biological, 40150-D001) or anti-MERS-CoV S (rabbit monoclonal, Sino Biological, 40069-R723) was added at a final concentration of 2 μg/mL and incubated at 37°C for 30 min. For nucleic acid staining, S-labled samples were diluted to 2×10^8^ particles/mL in PBS and incubated with SYTO9 at a ratio of 1:100 (v/v) at 37°C for 15 min prior to acquisition. Samples were analyzed using a Flow NanoAnalyzer (NanoFCM Inc., Xiamen, China) equipped with 488 nm and 638 nm lasers and three detection channels (side scatter [SSC], FITC, and PC5). Instrument calibration and data acquisition were performed according to the manufacturer’s instructions. Particles were gated based on size, and only events with diameters of 80–150 nm for pseudotyped particles or 60–140 nm for authentic virions were included. Data were analyzed using NanoFCM Profession software (v2.0).

### Immunoprecipitation and Pull-down assay

HEK293T cells were cultured in 10 cm dishes at a density of 2×10^6^ per dish for 12 h, and were transfected with plasmids (1 μg/mL) by Lipofectamine^TM^ 3000 Transfection Reagent (Thermo Fisher Scientific). 24 h later, cells were washed with PBS for 3 times and then lysed in M2 buffer (20 mM Tris-HCl pH 7.5, 0.5% Nonidet P-40, 250 mM NaCl, 3 mM EDTA, 3 mM EGTA) supplemented with complete protease inhibitor cocktail and PMSF, followed by centrifugation at 16,000 × g for 30 min at 4°C. The supernatants were incubated with ANTI-FLAG^®^M2 Affinity Gel (for FLAG IP), or with MagStrep “type3” XT beads (for Twin-Strep affinity purification), for 4 h at 4°C. The beads were washed 4 times with M2 buffer and boiled in 1×SDS-loading buffer for immunoblot analysis.

For Mass Spectrometry analysis, gel samples were processed by BIOMS Ltd. Peptides were separated on a nanoViper C18 column (75 μm × 250 mm, 2 μm) using an UltiMate 3000 UHPLC system coupled with an Orbitrap Exploris 480 mass spectrometer (Thermo Fisher Scientific) with a 60-minute gradient elution. Mass spectra were acquired in data-dependent acquisition (DDA) mode. Raw data were analyzed using MaxQuant (v2.0.1.0) against the Uniprot human protein database.

For ubiquitination assays, cells were lysed in RIPA buffer (20 mM Tris-HCl, pH 7.5; 150 mM NaCl; 1% Triton X-100; 10 mM EDTA; 1% sodium deoxycholate) supplemented with complete protease inhibitor cocktail, 1 mM PMSF, 1 mM DTT, PhosSTOP™ phosphatase inhibitor cocktail and 10 mM N-Ethylmaleimide (NEM).

Lysates were subjected to brief sonication on ice until visually clear, followed by centrifugation at 16,000 × g for 10 min at 4°C. Supernatants were adjusted to 1% final SDS concentration, boiled at 95°C for 5 min, then diluted tenfold with fresh RIPA buffer (containing 1 mM PMSF and 2 mM NEM) prior to incubation with ANTI-FLAG^®^ M2 Affinity Gel for 4 h at 4°C. Beads were washed 4 times with RIPA buffer and boiled in 1×SDS-loading buffer for immunoblot analysis.

### Periodic Acid-Schiff (PAS) Staining and Coomassie Brilliant Blue (CBB) Staining

Following SDS-PAGE, glycoproteins were visualized by PAS staining using a commercial kit (Real-Times, RTD6501) according to the manufacturer’s instructions. Briefly, gels were fixed in fixative solution, oxidized with periodic acid, incubated with Schiff’s reagent, and developed until distinct magenta bands appeared. For total protein detection, parallel gels were stained with InstantBlue Coomassie Protein Stain (Abcam, ab119211) for 30 min at room temperature, followed by destaining in distilled water until sharp, clear protein bands were resolved against a transparent background.

### Cell fractionation

HEK293T cells were cultured in 12-well plates at a density of 2×10^5^ per well for 12 h, and were transfected with plasmids (1 μg/well) by Lipofectamine^TM^ 3000 Transfection Reagent. 24 h later, cells were washed with PBS for 3 times and then lysed using the Minute™ Plasma Membrane/Protein Isolation and Cell Fractionation Kit (Invent Techbiologies, SM-005) according to the manufacturer’s instructions. Briefly, cells were homogenized by passage through the provided filtration column to generate native lysates with preserved organelle integrity. Following centrifugation at 700 × g for 1 min at 4°C, the pellet—enriched in nuclei and unlysed cells— was discarded, and the supernatant—containing plasma membrane fragments, intracellular membranes, and cytosolic components—was retained. This supernatant was then subjected to centrifugation at 16,000 × g for 30 min at 4°C; the resulting supernatant was designated as the cytosolic fraction, and the pellet as the total membrane fraction. The membrane pellet was resuspended directly in 1× SDS loading buffer, and the cytosolic fraction was mixed with 5 × SDS loading buffer. Both samples were then boiled for 5 min at 95°C for immunoblot analysis.

### LC-MS/MS analysis

Upper and lower bands were excised from SDS-PAGE gels, reduced with 5 mM DTT, and alkylated with 11 mM iodoacetamide. Gel pieces were digested with sequencing-grade modified trypsin overnight at 37°C. Peptides were extracted twice with 0.1% trifluoroacetic acid in 50% acetonitrile for 30 min and concentrated by vacuum centrifugation in a SpeedVac. Then, peptide mixtures were separated using a 60-min linear gradient at a flow rate of 300 nL/min on a Thermo-Dionex Ultimate 3000 HPLC system coupled to an Orbitrap Ascend mass spectrometer (Thermo Fisher Scientific). Peptides were loaded onto a self-packed fused silica analytical column (100 µm ID, 250 mm length; Upchurch) packed with C-18 resin (120 Å, 1.9 µm, Dr. Maisch GmbH). Mobile phase A consisted of 0.1% formic acid in water, and mobile phase B consisted of 80% acetonitrile with 0.1% formic acid. The mass spectrometer was operated in data-dependent acquisition mode using Xcalibur (v4.1). Full MS scans were acquired in the Orbitrap over a mass range of 300–1500 m/z at 120,000 resolutions, followed by top-speed MS/MS scans in the Orbitrap.

Raw data were analyzed using pGlyco 3.1^2,3^. Carbamidomethylation (C) was set as a fixed modification and methionine oxidation (M) as a variable modification. N-glycan compositions were searched against the built-in N-glycan database. Precursor and fragment mass tolerances were set to 20 ppm and 0.02 Da, respectively. Cleavage specificity was fully specific for trypsin with up to two missed cleavages allowed. Fragmentation was set to ETHCD. Peptide-, glycan-, and glycopeptide-level FDRs were controlled at 1%. Glycopeptide abundance was quantified using MS1 isotope envelope areas (IsotopeArea). Site-specific glycosylation intensity was calculated by summing glycopeptide intensities mapped to the same N-glycosylation site. Site intensities were normalized to the corresponding protein Intensity values obtained from MaxQuant (v2.7.5)^4,5^.

### Statistical analyses

Statistical analysis was performed using GraphPad Prism 9.0 software. All results are expressed as mean ± standard error of the mean (SEM). *P* values were determined by using unpaired two-tailed *t*-test. P values of < 0.05 were considered statistically significant between groups (ns, not significant; * *P* < 0.05; ** *P* < 0.01; *** *P* < 0.001; **** *P* < 0.0001).

## Data availability

Custom ImageJ/Fiji macros used for unbiased ROI selection are available on GitHub at https://github.com/chafinsondheimer945-pixel/random. The sequence of the SARS-CoV-2 JN.1 strain (SARS-CoV-2/human/CHN/IME-BJ202403/2024) has been deposited in the GenBase in National Genomics Data Center, Beijing Institute of Genomics, Chinese Academy of Sciences/China National Center for Bioinformation, under accession number C_AA160427.1 that is publicly accessible at https://ngdc.cncb.ac.cn/genbase^53,54^.

## Acknowledgements

This work was supported by grants from the National Key Research and Development Program of China (2022YFC2303700, 2021YFC2302400), National Natural Science Foundation of China (92469112, 92369114). We thank Yanjie Li for technical support during EM data collection. We thank the Tsinghua University Branch of China National Center for Protein Sciences (Beijing) for providing the cryo-EM facility support and thecomputational facility support. We thank Xiaolin Tian and Dr. Haiteng Deng in Center of Protein Analysis Technology, Tsinghua University, for MS analysis.

## Author contributions

Conceptualization, C.-F.Q. and Y.-J.H.; methodology, C.-F.Q., Y.-J.H. and L.L.; Investigation, Y.-J.H., L.L., W.R. H.Z., Y.-Q.D., Q.Y., L.Li and X.-Y.W.; writing—original draft, Y.-J.H. and L.L.; writing—review & editing, C.-F.Q.; funding acquisition, C.-F.Q., Y.-J.H., Y.-Q.D. and Q.C.; resources, C.-F.Q., Y.H., B.S., J.- J.Z. and J.Z.; supervision, C.-F.Q., J.Z. and Y.-J.H.

---

**Extended Data Fig. 1.**
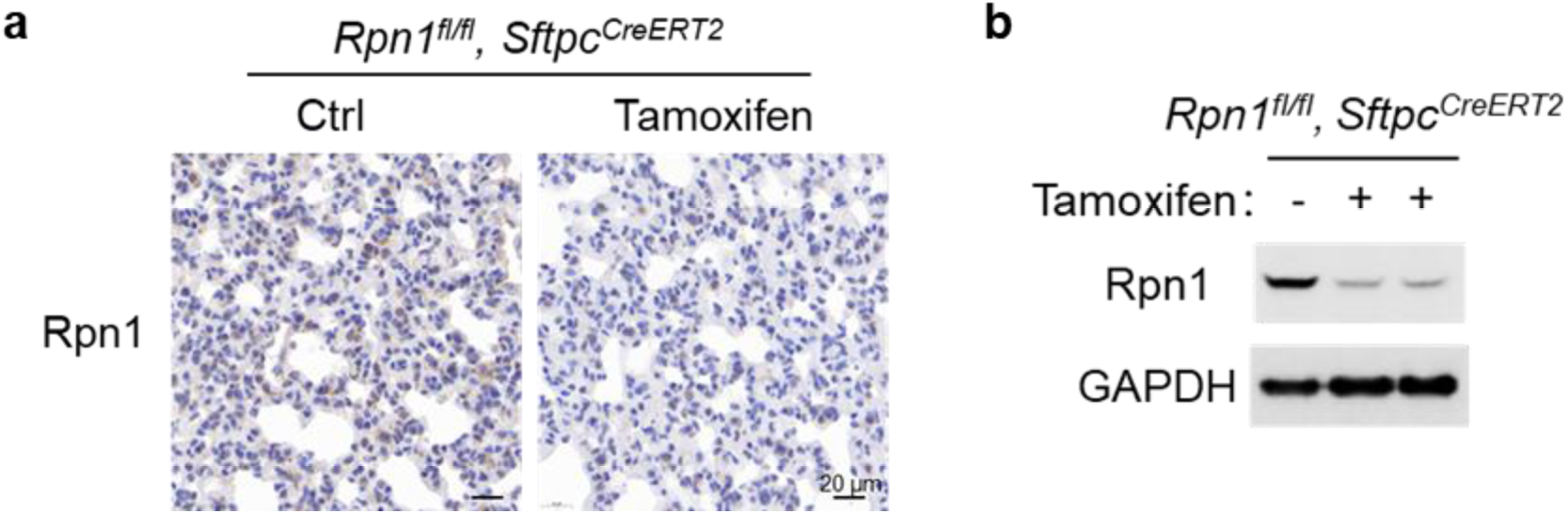
Validation of Rpn1 knockout efficiency in Rpn1-cKO Mice. **a**, Immunohistochemical staining of Rpn1 in lung sections from *Rpn1^fl/fl^, S*ftpc*^CreERT2^* mice treated with tamoxifen. **b**, Immunoblot analysis showing Rpn1 depletion in lungs of tamoxifen-treated *Rpn1^fl/fl^, S*ftpc*^CreERT2^* mice. GAPDH serves as a loading control.

**Extended Data Fig. 2.**
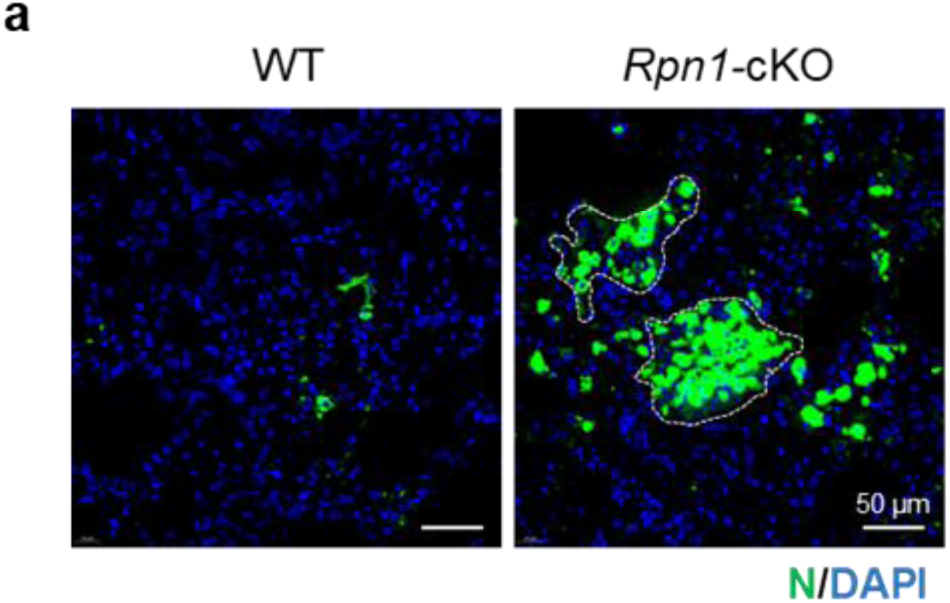
Enhanced SARS-CoV-2 infection in Rpn1-cKO mice. **a**, Immunofluorescence staining of lung sections from WT and *Rpn1-*cKO mice infected with SARS-CoV-2. Viral N protein (green) and nuclei (DAPI, blue) are shown. Outlined regions indicate multinucleated syncytial structures.

**Extended Data Fig. 3.**
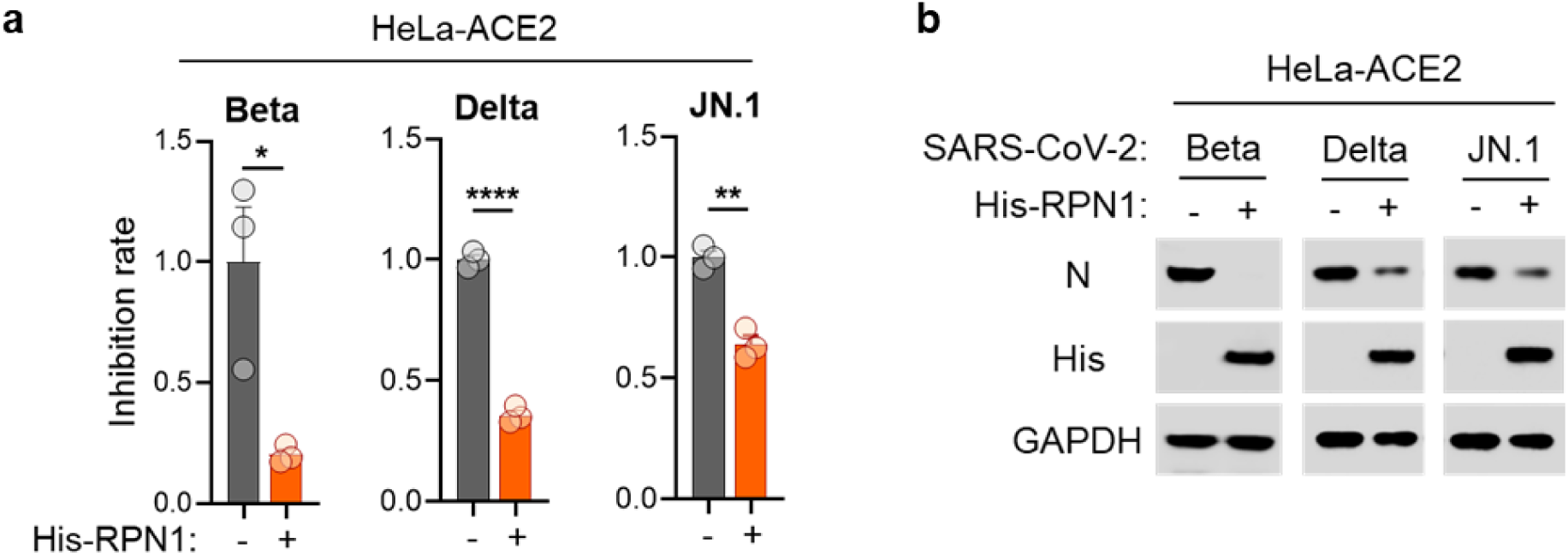
RPN1 inhibits infection by multiple SARS-CoV-2 variants. **a**,**b**, HeLa-ACE2 cells were transfected with *His-RPN1* for 24 h prior to infection with SARS-CoV-2 variants (Beta, Delta, and JN.1) at MOI=0.01 for 48 h. (**a**) Viral RNA levels in supernatants, quantified by qRT-PCR, are presented relative to the control group (set to 1) to calculate the percent inhibition conferred by RPN1 group. Data are presented as mean ± SEM of biological replicates (n = 3), two-tailed unpaired *t*-test. (* *P* < 0.05; ** *P* < 0.01; **** *P* < 0.0001) (**b**) Intracellular viral N protein and His-RPN1 expression were analyzed by immunoblotting. GAPDH serves as a loading control.

**Extended Data Fig. 4.**
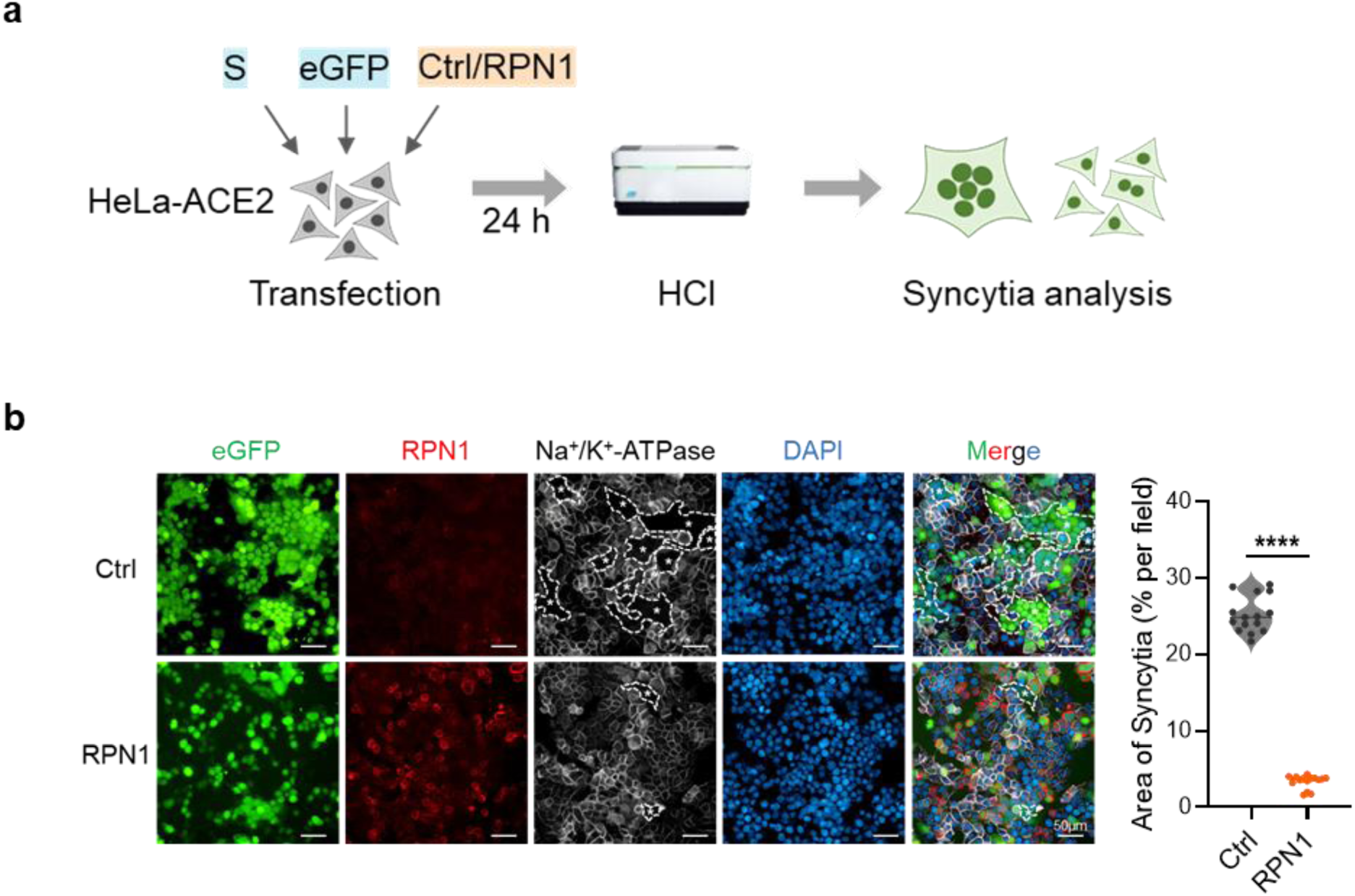
RPN1 inhibits the syncytium formation induced by the SARS-CoV-2 spike protein. **a**, Schematic diagram of the experimental workflow for S-mediated syncytia analysis. **b**, Immunofluorescence analysis of S-mediated syncytium formation in control or RPN1-overexpressing cells. eGFP (green) co-transfected with S was used to visualize S-mediated membrane fusion. RPN1 (red), Na⁺/K⁺-ATPase (membrane marker, gray), and nuclei (DAPI, blue) are shown. Dashed outlines with asterisks indicate representative syncytia. Quantification of syncytial area per field is shown on the right. Data are presented as mean ± SEM, two-tailed unpaired *t*-test. (**** *P* < 0.0001). Data represent at least three independent experiments.

**Extended Data Fig. 5.**
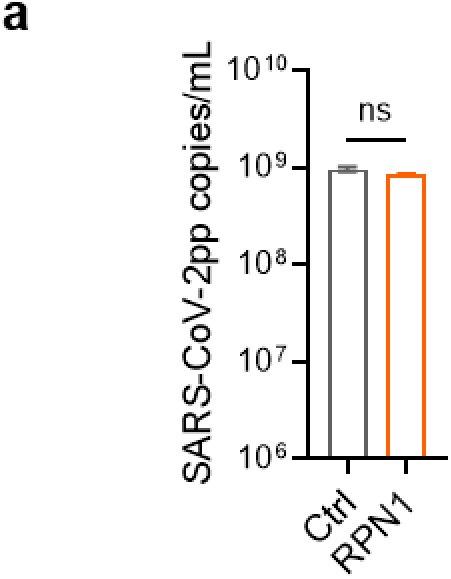
RPN1 does not affect the RNA level of pseudovirions. **a**, RNA levels of purified pseudotyped virus produced in control or RPN1-overexpressing cells, measured by qRT-PCR. Data are presented as mean ± SEM, two-tailed unpaired *t*-test. (ns, not significant). Data represent at least three independent experiments.

**Extended Data Fig. 6.**
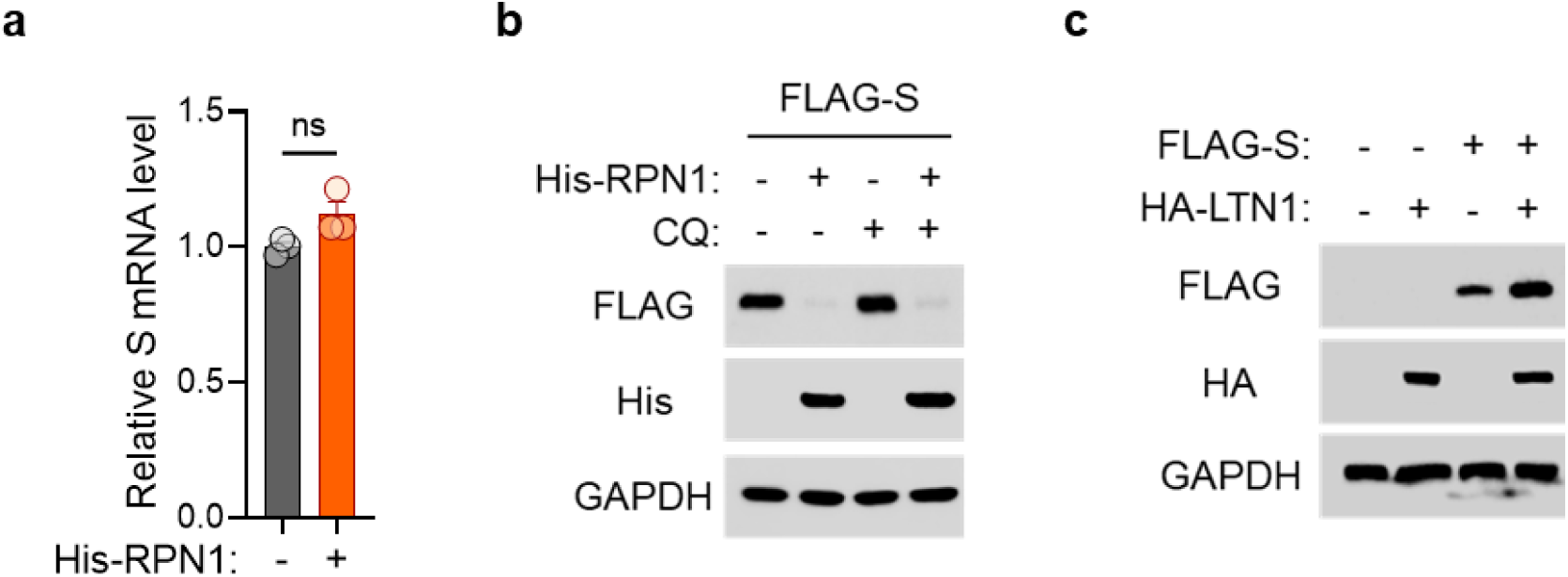
The spike protein is not downregulated by the lysosomal degradation pathway or the RQC pathway. **a**, *S* mRNA levels in cells co-expressing FLAG-S with or without His-RPN1, quantified by qRT-PCR and normalized to *GAPDH* mRNA. Values are presented relative to those of cells without RPN1. Data are presented as mean ± SEM of biological replicates (n = 3), two-tailed unpaired *t*-test. (ns, not significant). **b**, Immunoblot analysis of S protein in cells co-expressing FLAG-S and His-RPN1. Cells were transfected for 24 h followed by 50 μM CQ treatment for 12 h. **c**, Immunoblot analysis of S protein in cells co-expressing FLAG-S with or without HA-LTN1.

**Extended Data Fig. 7.**
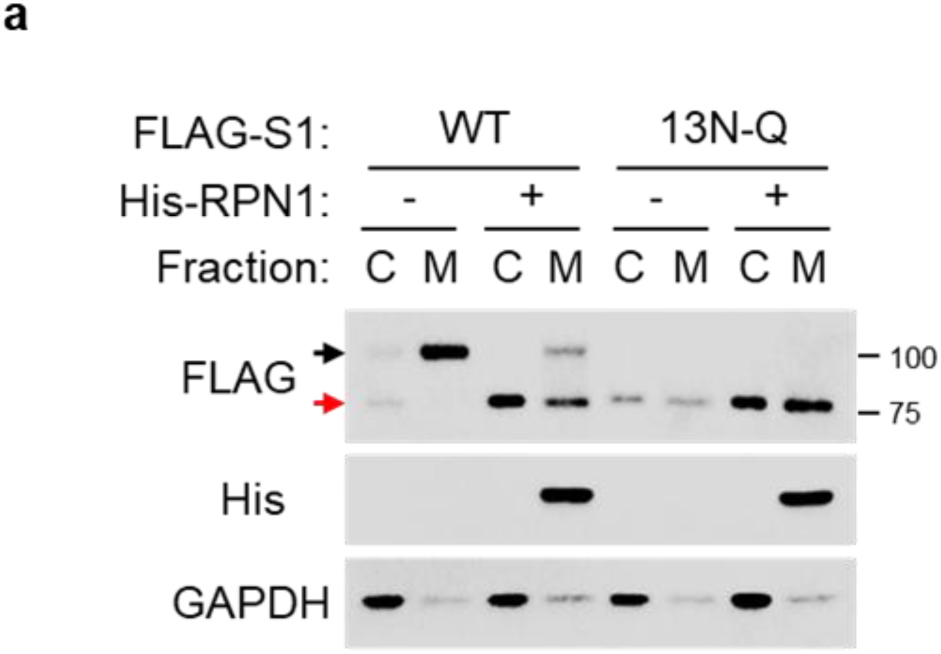
The lower band is the non-glycosylated form of spike protein. **a**, Immunoblot analysis of S protein in C and M fractions from cells expressing FLAG-S1 or mutant (13N-Q), with or without His-RPN1. GAPDH serves as a loading control.

**Extended Data Fig. 8.**
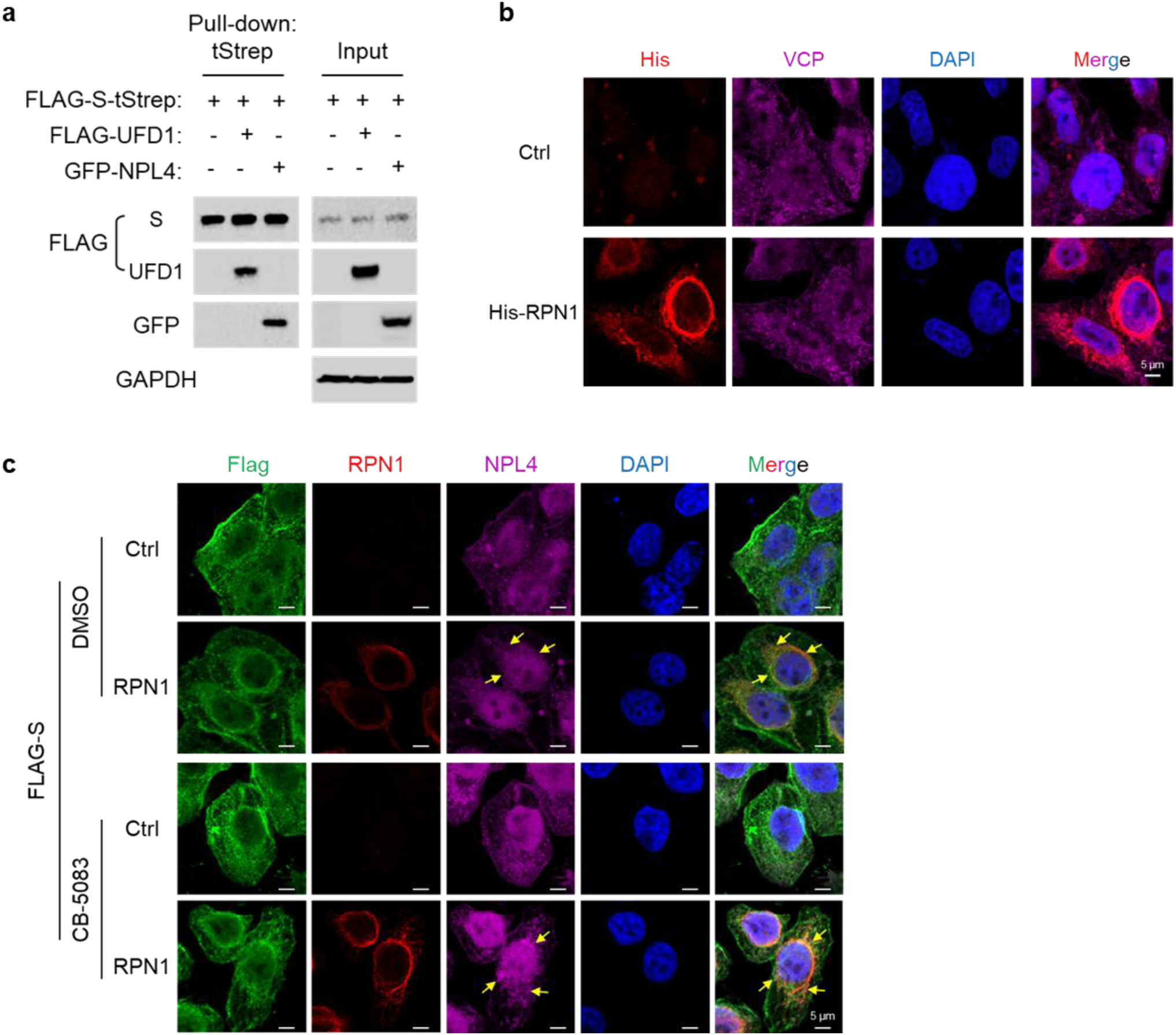
RPN1 induced the VCP/p97 complex-mediated ERAD of spike. **a**, C-terminally twin-strep (tStrep)-tagged FLAG-S was co-transfected with FLAG-UFD1 or GFP-NPL4 into HEK293T. tStrep affinity purification was performed, and co-precipitated proteins were probed with the indicated antibodies. GAPDH serves as a loading control. **b**, Control/His-RPN1 were transfected into the HEK293T for 24 h. Immunofluorescence staining shows RPN1 (red), VCP (magenta), and nuclei (DAPI, blue). **c**, FLAG-S and control/His-RPN1 were co-transfected into the HEK293T for 24 h, followed by treatment with DMSO or 1 μM CB-5083 for 12 h. Immunofluorescence staining shows S (FLAG, green), RPN1 (red), NPL4 (magenta), and nuclei (DAPI, blue).

**Extended Data Fig. 9.**
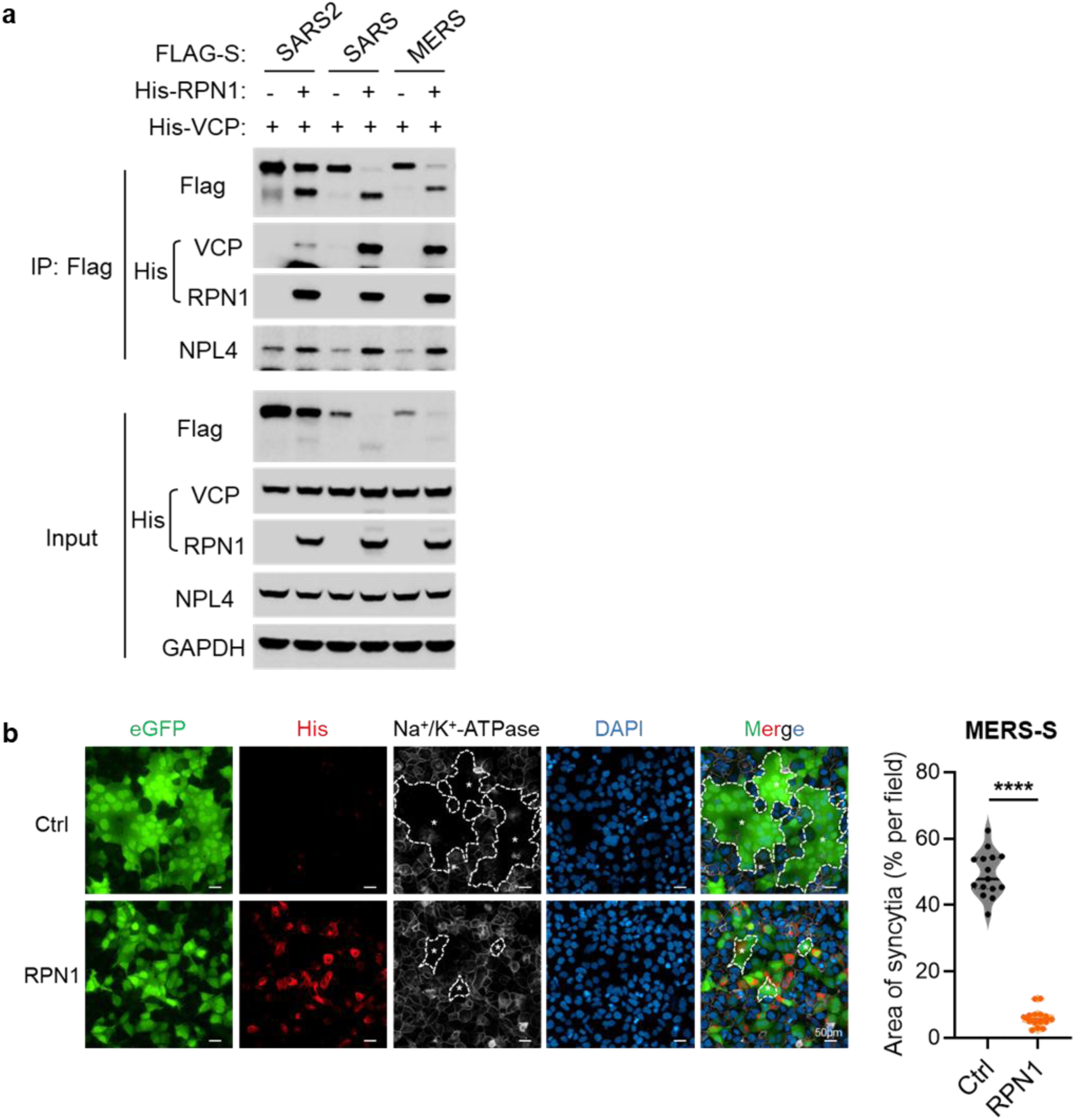
RPN1-mediated spike ERAD is conserved across highly pathogenic coronaviruses. **a**, FLAG-S (SARS2, SARS or MERS) and control/His-RPN1 were co-transfected into the HEK293T overexpressing His-VCP. FLAG immunoprecipitation was performed, and co-precipitated proteins were probed with the indicated antibodies. GAPDH serves as a loading control. **b**, Immunofluorescence analysis of MERS-S-mediated syncytium formation in Huh7 cells transfected with control or RPN1. eGFP (green) was used to visualize S-mediated membrane fusion. RPN1 (His, red), Na⁺/K⁺-ATPase (membrane marker, gray), and nuclei (DAPI, blue) are shown. Dashed outlines with asterisks indicate representative syncytia. Quantification of total syncytial area per field is presented on the right. Data are presented as mean ± SEM, two-tailed unpaired *t*-test. (**** *P* < 0.0001). Data represent at least three independent experiments.

**Extended Data Fig. 10.**
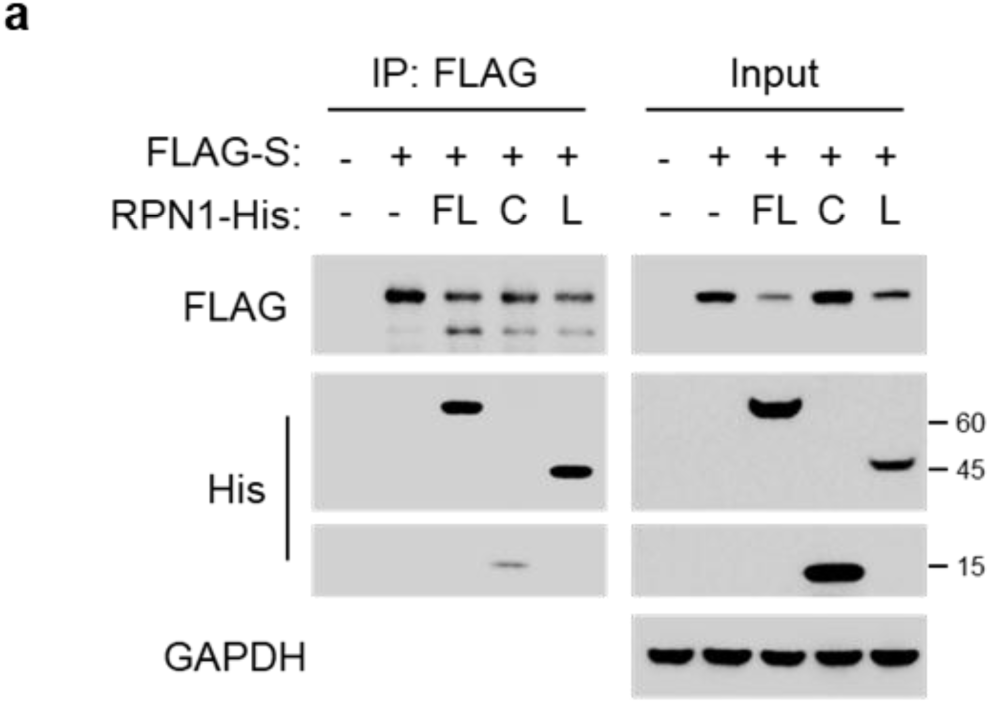
The luminal domain of RPN1 interacts directly with the SARS-CoV-2 spike protein. **a**, FLAG-S and control/His-RPN1 (FL, C or L) were co-transfected into the HEK293T cells. FLAG immunoprecipitation was performed, and co-precipitated RPN1 proteins were probed with His antibody. GAPDH serves as a loading control.

## References

1 V’Kovski, P., Kratzel, A., Steiner, S., Stalder, H. & Thiel, V. Coronavirus biology and replication: implications for SARS-CoV-2. Nat Rev Microbiol 19, 155–170 (2021). 10.1038/s41579-020-00468-6

2 Lu, L., Su, S., Yang, H. & Jiang, S. Antivirals with common targets against highly pathogenic viruses. Cell 184, 1604–1620 (2021). 10.1016/j.cell.2021.02.013

3 Rebendenne, A. et al. Bidirectional genome-wide CRISPR screens reveal host factors regulating SARS-CoV-2, MERS-CoV and seasonal HCoVs. Nat Genet 54, 1090–1102 (2022). 10.1038/s41588-022-01110-2

4 Schmidt, N. et al. The SARS-CoV-2 RNA-protein interactome in infected human cells. Nat Microbiol 6, 339–353 (2021). 10.1038/s41564-020-00846-z

5 Baggen, J., Vanstreels, E., Jansen, S. & Daelemans, D. Cellular host factors for SARS-CoV-2 infection. Nat Microbiol 6, 1219–1232 (2021). 10.1038/s41564-021-00958-0

6 Chen, B., Farzan, M. & Choe, H. SARS-CoV-2 spike protein: structure, viral entry and variants. Nat Rev Microbiol 23, 455–468 (2025). 10.1038/s41579-025-01185-8

7 Cui, J., Li, F. & Shi, Z. L. Origin and evolution of pathogenic coronaviruses. Nat Rev Microbiol 17, 181–192 (2019). 10.1038/s41579-018-0118-9

8 Minigulov, N., Boranbayev, K., Bekbossynova, A., Gadilgereyeva, B. & Filchakova, O. Structural proteins of human coronaviruses: what makes them different? Front Cell Infect Microbiol 14, 1458383 (2024). 10.3389/fcimb.2024.1458383

9 Uraki, R., Korber, B., Diamond, M. S. & Kawaoka, Y. SARS-CoV-2 variants: biology, pathogenicity, immunity and control. Nat Rev Microbiol 24, 8–28 (2026). 10.1038/s41579-025-01255-x

10 Jackson, C. B., Farzan, M., Chen, B. & Choe, H. Mechanisms of SARS-CoV-2 entry into cells. Nat Rev Mol Cell Biol 23, 3–20 (2022). 10.1038/s41580-021-00418-x

11 Yao, H. et al. Molecular Architecture of the SARS-CoV-2 Virus. Cell 183, 730–738 e713 (2020). 10.1016/j.cell.2020.09.018

12 Letko, M., Marzi, A. & Munster, V. Functional assessment of cell entry and receptor usage for SARS-CoV-2 and other lineage B betacoronaviruses. Nat Microbiol 5, 562–569 (2020). 10.1038/s41564-020-0688-y

13 Braga, L. et al. Drugs that inhibit TMEM16 proteins block SARS-CoV-2 spike-induced syncytia. Nature 594, 88–93 (2021). 10.1038/s41586-021-03491-6

14 Palchevska, O. & Dominguez, F. Syncytium: the viral escape room secret to persistent infection of SARS-CoV-2. Front Microbiol 16, 1561274 (2025). 10.3389/fmicb.2025.1561274

15 Peluso, M. J. & Deeks, S. G. Mechanisms of long COVID and the path toward therapeutics. Cell 187, 5500–5529 (2024). 10.1016/j.cell.2024.07.054

16 Ke, Z. et al. Structures and distributions of SARS-CoV-2 spike proteins on intact virions. Nature 588, 498–502 (2020). 10.1038/s41586-020-2665-2

17 Zhang, L. et al. SARS-CoV-2 spike-protein D614G mutation increases virion spike density and infectivity. Nat Commun 11, 6013 (2020). 10.1038/s41467-020-19808-4

18 Zhang, F. et al. SARS-CoV-2 spike glycosylation affects function and neutralization sensitivity. mBio 15, e0167223 (2024). 10.1128/mbio.01672-23

19 Lusvarghi, S. et al. Effects of N-glycan modifications on spike expression, virus infectivity, and neutralization sensitivity in ancestral compared to Omicron SARS-CoV-2 variants. PLoS Pathog 19, e1011788 (2023). 10.1371/journal.ppat.1011788

20 Li, Y. et al. A substitution at the cytoplasmic tail of the spike protein enhances SARS-CoV-2 infectivity and immunogenicity. EBioMedicine 110, 105437 (2024). 10.1016/j.ebiom.2024.105437

21 Guay, K. P., Chou, W. C., Canniff, N. P., Paul, K. B. & Hebert, D. N. N-glycan-dependent protein maturation and quality control in the ER. Nat Rev Mol Cell Biol 26, 926–939 (2025). 10.1038/s41580-025-00855-y

22 Gemmer, M. et al. Visualization of translation and protein biogenesis at the ER membrane. Nature 614, 160–167 (2023). 10.1038/s41586-022-05638-5

23 Wilson, C. M., Roebuck, Q. & High, S. Ribophorin I regulates substrate delivery to the oligosaccharyltransferase core. Proc Natl Acad Sci U S A 105, 9534–9539 (2008). 10.1073/pnas.0711846105

24 Ramirez, A. S., Kowal, J. & Locher, K. P. Cryo-electron microscopy structures of human oligosaccharyltransferase complexes OST-A and OST-B. Science 366, 1372–1375 (2019). 10.1126/science.aaz3505

25 Liang, J. R. et al. A Genome-wide ER-phagy Screen Highlights Key Roles of Mitochondrial Metabolism and ER-Resident UFMylation. Cell 180, 1160–1177 e1120 (2020). 10.1016/j.cell.2020.02.017

26 Kucej, M., Fermaintt, C. S., Yang, K., Irizarry-Caro, R. A. & Yan, N. Mitotic Phosphorylation of TREX1 C Terminus Disrupts TREX1 Regulation of the Oligosaccharyltransferase Complex. Cell Rep 18, 2600–2607 (2017). 10.1016/j.celrep.2017.02.051

27 Hwang, H. J. et al. Therapy-induced senescent cancer cells contribute to cancer progression by promoting ribophorin 1-dependent PD-L1 upregulation. Nat Commun 16, 353 (2025). 10.1038/s41467-024-54132-1

28 Huang, Y. J. et al. Identification of oligosaccharyltransferase as a host target for inhibition of SARS-CoV-2 and its variants. Cell Discov 7, 116 (2021). 10.1038/s41421-021-00354-2

29 Sanders, D. W. et al. SARS-CoV-2 requires cholesterol for viral entry and pathological syncytia formation. Elife 10 (2021). 10.7554/eLife.65962

30 Baggen, J. et al. TMEM106B is a receptor mediating ACE2-independent SARS-CoV-2 cell entry. Cell 186, 3427–3442 e3422 (2023). 10.1016/j.cell.2023.06.005

31 Li, F. Structure, Function, and Evolution of Coronavirus Spike Proteins. Annu Rev Virol 3, 237–261 (2016). 10.1146/annurev-virology-110615-042301

32 Gao, Y., Zhu, Y., Sun, Q. & Chen, D. Argonaute-dependent ribosome-associated protein quality control. Trends Cell Biol 33, 260–272 (2023). 10.1016/j.tcb.2022.07.007

33 Lemberg, M. K. & Strisovsky, K. Maintenance of organellar protein homeostasis by ER-associated degradation and related mechanisms. Mol Cell 81, 2507–2519 (2021). 10.1016/j.molcel.2021.05.004

34 Christianson, J. C., Jarosch, E. & Sommer, T. Mechanisms of substrate processing during ER-associated protein degradation. Nat Rev Mol Cell Biol 24, 777–796 (2023). 10.1038/s41580-023-00633-8

35 Watanabe, Y., Allen, J. D., Wrapp, D., McLellan, J. S. & Crispin, M. Site-specific glycan analysis of the SARS-CoV-2 spike. Science 369, 330–333 (2020). 10.1126/science.abb9983

36 Wang, H. H., Biunno, I., Sun, S. & Qi, L. SEL1L-HRD1-mediated ERAD in mammals. Nat Cell Biol 27, 1063–1073 (2025). 10.1038/s41556-025-01690-1

37 Olzmann, J. A., Kopito, R. R. & Christianson, J. C. The mammalian endoplasmic reticulum-associated degradation system. Cold Spring Harb Perspect Biol 5 (2013). 10.1101/cshperspect.a013185

38 Braxton, J. R. & Southworth, D. R. Structural insights of the p97/VCP AAA+ ATPase: How adapter interactions coordinate diverse cellular functionality. J Biol Chem 299, 105182 (2023). 10.1016/j.jbc.2023.105182

39 Pontifex, C. S., Zaman, M., Fanganiello, R. D., Shutt, T. E. & Pfeffer, G. Valosin-Containing Protein (VCP): A Review of Its Diverse Molecular Functions and Clinical Phenotypes. Int J Mol Sci 25 (2024). 10.3390/ijms25115633

40 Zhou, H. J. et al. Discovery of a First-in-Class, Potent, Selective, and Orally Bioavailable Inhibitor of the p97 AAA ATPase (CB-5083). J Med Chem 58, 9480–9497 (2015). 10.1021/acs.jmedchem.5b01346

41 Liu, Y. et al. USP13 antagonizes gp78 to maintain functionality of a chaperone in ER-associated degradation. Elife 3, e01369 (2014). 10.7554/eLife.01369

42 Raj, V. S. et al. Dipeptidyl peptidase 4 is a functional receptor for the emerging human coronavirus-EMC. Nature 495, 251–254 (2013). 10.1038/nature12005

43 Qian, Z., Dominguez, S. R. & Holmes, K. V. Role of the spike glycoprotein of human Middle East respiratory syndrome coronavirus (MERS-CoV) in virus entry and syncytia formation. PLoS One 8, e76469 (2013). 10.1371/journal.pone.0076469

44 Simmons, G. et al. Characterization of severe acute respiratory syndrome-associated coronavirus (SARS-CoV) spike glycoprotein-mediated viral entry. Proc Natl Acad Sci U S A 101, 4240–4245 (2004). 10.1073/pnas.0306446101

45 Marceau, C. D. et al. Genetic dissection of Flaviviridae host factors through genome-scale CRISPR screens. Nature 535, 159–163 (2016). 10.1038/nature18631

46 Bagchi, P. Endoplasmic reticulum in viral infection. Int Rev Cell Mol Biol 350, 265–284 (2020). 10.1016/bs.ircmb.2019.10.005

47 Liu, S. R. et al. Coronaviral nsp6 hijacks ERAD machinery to facilitate lipolysis and supply membrane components for DMV growth. Nat Commun 16, 10094 (2025). 10.1038/s41467-025-65118-y

48 Choy, M. M. et al. A Non-structural 1 Protein G53D Substitution Attenuates a Clinically Tested Live Dengue Vaccine. Cell Rep 31, 107617 (2020). 10.1016/j.celrep.2020.107617

49 Zeng, W. F., Cao, W. Q., Liu, M. Q., He, S. M. & Yang, P. Y. Precise, fast and comprehensive analysis of intact glycopeptides and modified glycans with pGlyco3. Nat Methods 18, 1515–1523 (2021). 10.1038/s41592-021-01306-0

50 Kong, S. et al. pGlycoQuant with a deep residual network for quantitative glycoproteomics at intact glycopeptide level. Nat Commun 13, 7539 (2022). 10.1038/s41467-022-35172-x

51 Peng, H., Wang, H., Kong, W., Li, J. & Goh, W. W. B. Optimizing differential expression analysis for proteomics data via high-performing rules and ensemble inference. Nat Commun 15, 3922 (2024). 10.1038/s41467-024-47899-w

52 Cox, J. & Mann, M. MaxQuant enables high peptide identification rates, individualized p.p.b.-range mass accuracies and proteome-wide protein quantification. Nat Biotechnol 26, 1367–1372 (2008). 10.1038/nbt.1511

53 Congfan, B. et al. GenBase: A Nucleotide Sequence Database. Genomics Proteomics Bioinformatics 22, qzae047 (2024). 10.1093/gpbjnl/qzae047

54 Partners, C.-N. M. a. Database Resources of the National Genomics Data Center, China National Center for Bioinformation in 2025. Nucleic Acids Res 53, D30-D44 (2024). 10.1093/nar/gkae978

